# De novo protein synthesis in distinct centrolateral amygdala interneurons is required for associative emotional memories

**DOI:** 10.1101/2020.04.08.028233

**Authors:** Prerana Shrestha, Zhe Shan, Maggie Marmarcz, Karen San Agustin Ruiz, Adam Taye Zerihoun, Chien-Yu Juan, Pedro Manuel Herrero-Vidal, Jerry Pelletier, Nathaniel Heintz, Eric Klann

## Abstract

To survive in a dynamic environment, animals need to identify and appropriately respond to stimuli that signal danger^1,2^. At the same time, animal survival also depends on suppressing the threat response during a stimulus that predicts absence of threat, i.e. safety^3-5^. Understanding the biological substrates of differential threat memories in which animals learn to flexibly switch between expressing and suppressing defensive responses to a threat-predictive cue and a safety cue, respectively, is critical for developing treatments for memory disorders such as PTSD^6^. A key brain area for processing and storing threat memories is the centrolateral amygdala (CeL), which receives convergent sensory inputs from the parabrachial nucleus and the basolateral amygdala and connects directly to the output nucleus of amygdala, the centromedial nucleus, to mediate defensive responses^7-9^. Despite a plethora of studies on the importance of neuronal activity in specific CeL neuronal populations during memory acquisition and retrieval^10-12^, little is known about regulation of their protein synthesis machinery. Consolidation of long-term, but not short-term, threat memories requires de novo protein synthesis, which suggests that the translation machinery in CeL interneurons is tightly regulated in order to stabilize associative memories. Herein, we have applied intersectional chemogenetic strategies in CeL interneurons to block cell type-specific translation initiation programs that are sensitive to depletion of eukaryotic initiation factor 4E (eIF4E) and phosphorylation of eukaryotic initiation factor 2α (p-eIF2α), respectively. We show that in a differential threat conditioning paradigm, de novo translation in somatostatin-expressing (SOM) interneurons in the CeL is necessary for long-term storage of conditioned threat response whereas de novo translation in protein kinase Cδ-expressing (PKCδ) interneurons in the CeL is essential for storing conditioned response inhibition to a safety cue. Further, we show that oxytocinergic neuromodulation of PKCδ interneurons during differential threat learning is important for long-lasting cued threat discrimination. Our results indicate that the molecular elements of a differential threat memory trace are compartmentalized in distinct CeL interneuron populations and provide new mechanistic insight into the role of de novo protein synthesis in consolidation of long-term memories.

---

Neurons have evolved to both respond dynamically to their environment at millisecond time scales and yet can store information stably for a much longer period of time. The latter mode of stabilizing information in mnemonic processes requires *de novo* translation^13,14^. Tight regulation of translation occurs during initiation where the two major rate-limiting steps are the assembly of the eIF2-tRNA_i_^Met^ ternary complex and the ^m7^GpppN cap-binding complex^15^. Bidirectional control of protein synthesis can be mediated by altering the levels of these two complexes. As part of the integrated stress response, eIF2α kinases phosphorylate eIF2α and this in turn inhibits the eIF2 guanine exchange factor eIF2B, effectively blocking recycling of the ternary complex to shutdown general translation. On the other hand, dephosphorylation of eIF2α occurs following memory formation, allowing the requisite *de novo* translation to initiate^16^. Likewise, the formation of the ^m7^GpppN cap-binding complex is essential for cap-dependent translation initiation. Central to the regulation of cap-dependent translation is the mammalian target of rapamycin complex I (mTORC1) signaling pathway. Activation of mTORC1 triggers initiation of cap-dependent translation via phosphorylation of eukaryotic initiation factor 4E (eIF4E)-binding proteins (4EBPs) and p70 S6 kinase 1 (S6K1). Phosphorylation of 4E-BPs results in the release of eIF4E, which then becomes incorporated into the eIF4F complex, along with the modular scaffolding protein eIF4G and the RNA helicase eIF4A to initiate cap-dependent translation. Phosphorylation of S6K1 leads to phosphorylation of downstream targets including ribosomal protein S6, eIF4B, and PDCD4, an event which promotes translation^15, 17^. Although both eIF2 and mTORC1 pathways regulate key steps in translation initiation, these are generally viewed as separate translation control pathways with largely non-overlapping molecular outcomes^18-21^.

We developed a differential threat conditioning paradigm using interleaved presentations of a shock-predictive tone (paired conditioned stimulus, CS+) that co-terminated with a footshock (unconditioned stimulus, US) and a safety-predictive tone that predicted absence of the footshock (CS-) within session (Fig. 1a and b). The Box-Only control group was placed in the training context but did not get exposed to either CS+ or CS-. The Unpaired training group (Fig. 1b) was exposed to all three stimuli (CS+, CS- and US) in scrambled order, precluding any tone-shock contingency. Compared to the Unpaired group, mice in the Paired training group learned the CS+-US association during training with progressive increase in freezing to successive CS presentations (Fig. 1c, Extended Data Fig. 1a) even though both groups increased freezing behavior post-tone (Extended Data Fig. 1b). When the mice were tested for long-term memory (LTM), paired training resulted in mice exhibiting a high freezing response to the CS+ while suppressing the response to CS- (Fig.1c-d), with a robust discrimination index outcome compared to Box-Only and Unpaired controls (Fig. 1e, Extended Data Fig. 1c). Notably, the freezing response to CS- was higher than the negligible freezing behavior during pre-CS period (Extended Data Fig.1d-e). Increasing the number of CS+-US pairings from 3 to 5 increased freezing with successive CS presentations during memory acquisition (Extended Data Fig. 1f), but did not improve the freezing response to either the CS+ and CS- or the discrimination index during LTM (Extended Data Fig. 1g-h), indicating that the learned behavior had reached an asymptote after 3 pairings. Biochemical analysis of the amygdala showed that activation of mTORC1, as indicated by phosphorylation of S6K1, occurs in Paired animals but not in Box-Only or Unpaired groups (Fig. 1f). Notably, dephosphorylation of eIF2α occurred in the differentially threat conditioned group as well as the Unpaired group, indicating pathway divergence of translation programs in capturing the shock experience versus tone-shock contingencies (Fig. 1f). We next focused on SOM and PKCδ interneuron (IN) subpopulations^22-25^ that each constitute approximately half of all neurons in the CeL (Extended Data Fig. 2a-b) and are largely distinct (Fig. 1g, Extended Data Fig. 2c-d). We found that phosphorylation of ribosomal protein S6 at Ser 235/6 was significantly increased in both SOM and PKCδ INs in the Paired group compared to Box-Only and Unpaired controls, indicating activation of the mTORC1 pathway by differential threat conditioning (Extended Data Fig. 2f-g). We then utilized *in vivo* surface sensing of translation (SUnSET) to label newly synthesized proteins with the synthetic tyrosyl-tRNA analog puromycin in awake behaving mice. A significant increase in *de novo* translation in CeL, specifically in PKCδ INs, was observed in the Paired group compared to both Unpaired and Box-only controls (Fig. 1h).

**Figure 1.**
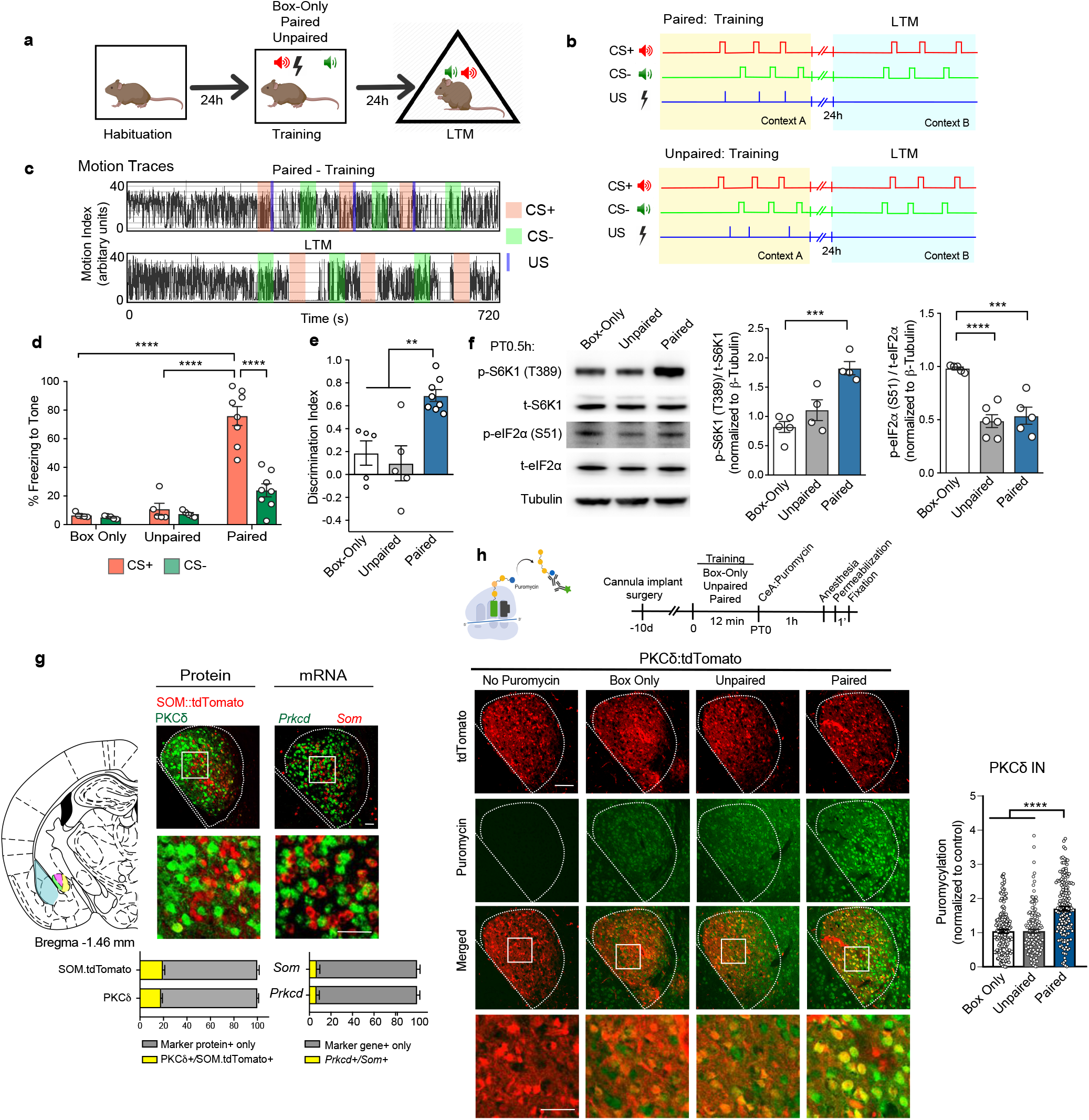
Differential threat conditioningd promotes *de novo* translation in CeL INs. a) Behavior scheme for differential cued threat conditioning. b) Schematic of the behavior protocol for the Paired training group (top) and the Unpaired training group (bottom). c) Representative motion traces for the Paired group during training and LTM test. CS+ in orange block, CS- in green block and US in violet block. d) During LTM, the Paired group displayed a robust freezing response to CS+ compared to Box-Only and Unpaired groups. Interaction (Training X CS): F(2,30)=20.41, p<0.0001; Training: F(2,30)=60.08, p<0.0001, CS: F(1,30)=22.86, p<0.0001. p=5-8/group. e) The Paired group exhibited a high discrimination index for cued threat compared to controls. F(2,15)=12.01, p=0.0008. f) Representative immunoblots for mTORC1 and eIF2 pathway indicators: p-S6K1 (T389), t-S6K1, p-eIF2α (S51), t-eIF2α and ß-Tubulin (left). g) p-S6K1 (T389) was significantly elevated in amygdala lysate of the Paired group compared to the Box-Only control (left) whereas dephosphorylation of eIF2α (S51) occurred in both Unpaired and Paired groups (right). p-S6K1: F(2,10)=16.41, p=0.0007. n=4-5/group. p-eIF2α: F(2,13)=20.94, p<0.0001. n=4-6/group. g) Schematic of the coronal brain section, Bregma −1.46 mm with amygdala subnuclei (left). Immunostaining for PKCδ in SOM tdTomato mice revealed largely distinct neuronal populations expressing SOM and PKCδ gene driver (middle). 18.06% of PKCδ+ neurons co-expressed SOM.tdTomato whereas 19.56% of SOM.tdTomato neurons co-expressed PKCδ. n=3/group. Small molecule fluorescent *in situ* hybridization (smFISH) for *Prkcd* and *Som* RNAs reveal discrete neuronal subpopulations in CeL (right) with double positive cells representing 6.63% of *Prkcd+* cells and 6.94% of *Som+* cells. n = 3/ group. h) Schematic for the *in vivo* de novo translation labeling assay with puromycin infusion in central amygdala (top). *De novo* translation was significantly upregulated in PKCδ INs in the Paired training group compared to Box-Only and Unpaired controls (bottom). Insets show higher magnification. F(2,482)=44.18, p<0.0001. n=158-165/group. Statistical tests: Two-way ANOVA with Bonferroni’s post-hoc test (d), One way ANOVA with Bonferroni’s post-hoc test (e, f, g, i, j). Data are presented as mean +SEM. *p<0.05, **p<0.01, ***p<0.001, ****p<0.0001. n.s. nonsignificant. Scale bar, 50 μm.

**Figure 2.**
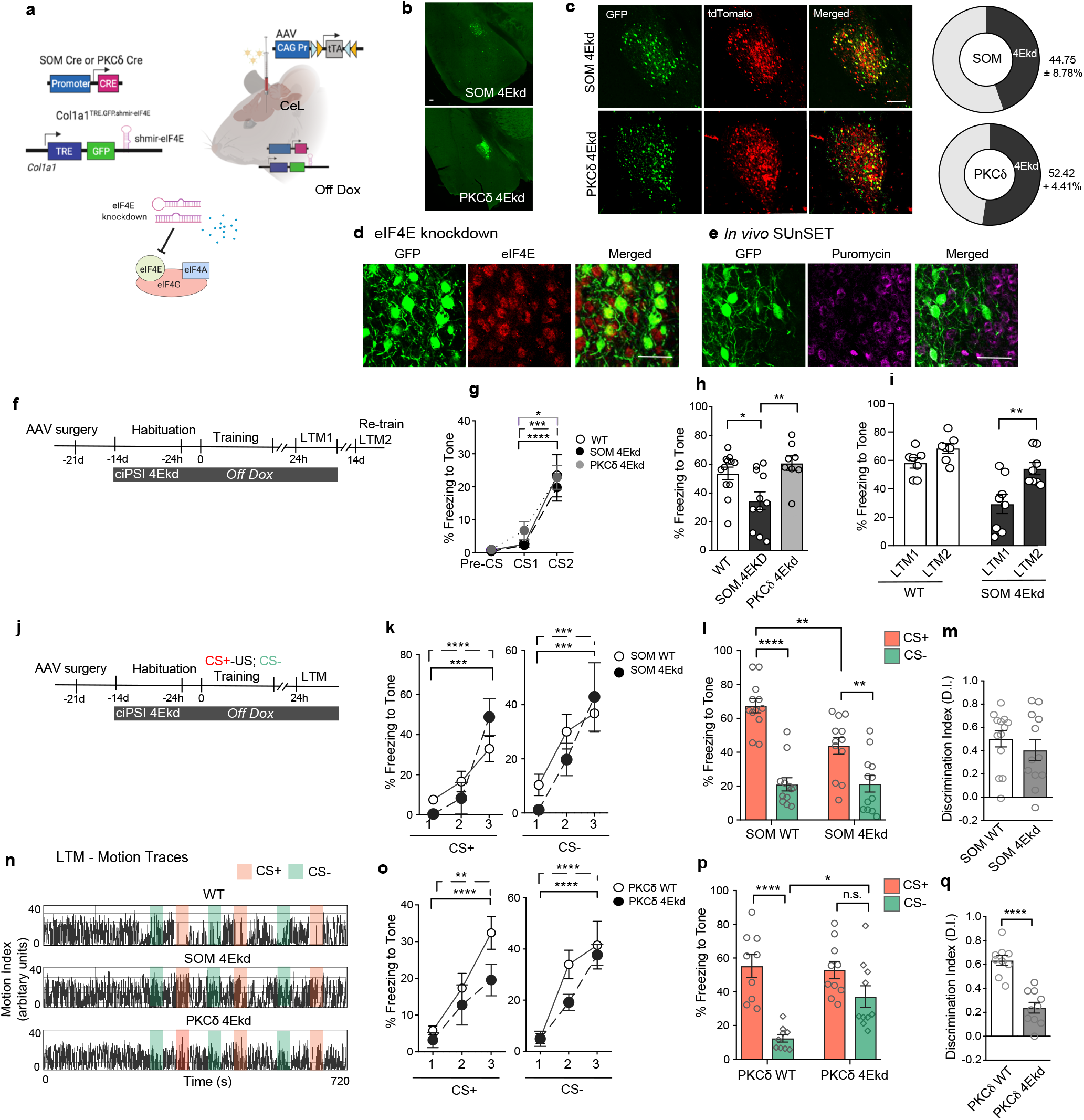
Cell type-specific inhibition of cap-dependent translation in CeL interneurons. a) Intersectional chemogenetic strategy for knocking down eIF4E in CeL SOM and PKCδ neurons. b) Viral delivery of AAV.CAG Pr.DIO.tTA in the CeL of SOM/ PKCδ.TRE GFP.shmir-eIF4E animals mediates cre-dependent expression of GFP and shmiR-eIF4E in SOM and PKCδ INs. c) Proportion of endogenous SOM and PKCδ INs targeted using the chemogenetic strategy. n=3/group d) GFP.shmiR-eIF4E knocks down eIF4E, and e) subsequently inhibits *de novo* translation as measured using *in vivo* puromycylation. f) Behavior paradigm for simple cued threat conditioning. g) XY plot showing simple threat conditioning in WT, SOM 4Ekd and PKCδ 4Ekd groups. CS: F(2,56)=31.88, p<0.0001. n= 8-12/group. h) SOM 4Ekd mice display significantly impaired LTM compared to both WT and PKCδ 4Ekd mice. F(2,28)=6.41, p=0.0051. n=8-12/group. i) Re-training the SOM 4Ekd mice after placing on Dox diet for 14 days completely rescues the memory deficit. Drug: F(1,13)=12.33, p=0.0038; Genotype: F(1,13)=21.13, p=0.0005. SOM 4Ekd: LTM1 vs LTM2, p=0.006. n=7-8/group. j) Behavior paradigm for cued differential threat conditioning. k) Normal memory acquisition in SOM 4Ekd mice as they show a progressive increase in freezing response to successive presentations of the CS. CS+: F(2,26)=34.66, p<0.0001. CS-: F(2,26)=20.81, p<0.0001. l) SOM 4Ekd mice have a significant impairment in CS+ threat LTM compared to SOM WT control, despite showing an equivalent safety response to CS-. Interaction (Genotype X CS): F(1,44) =7.079, p=0.0108. Genotype: F(1,44)=6.68, p=0.013. n=12/group. m) Normal cue discrimination index for SOM 4EKd mice compared to controls. n) Representative motion traces during differential threat LTM test for WT, SOM 4Ekd and PKCδ 4Ekd animals. o) Normal memory acquisition in PKCδ 4Ekd mice compared to PKCδ WT controls. CS+: F(2,34)=24.67, p<0.0001; CS-: F(2,34)=36.84, p<0.0001. n=9-10/group. p) PKCδ 4Ekd mice display a significant impairment in safety LTM to CS- despite showing comparable threat LTM to CS+. Interaction (Genotype X CS): F(1,34)=6.19, p=0.0179. Genotype: F(1,34)=4.17, p=0.049. CS: F(1,34)=28.60, p<0.0001. n=9-10/group. q) Discrimination index for cued threat is significantly impaired in PKCδ 4Ekd group compared with PKCδ WT controls. p<0.0001. n=9-10/group. Statistical tests: RM Oneway ANOVA (g, k, o), One-way ANOVA with Bonferroni’s post-hoc test (h), RM Two-way ANOVA with Bonferroni’s post-hoc test (i), Two-way ANOVA with Bonferroni’s post-hoc test (l, p) and Unpaired t-test (m, q). Data are presented as mean +SEM. *p<0.05, **p<0.01, ***p<0.001, ****p<0.0001. n.s. nonsignificant. Scale bar, 100 μm (b,c) and 50 μm (d,e).

To establish a causal role for cap-dependent translation in CeL interneurons in differential threat memories, we devised an intersectional chemogenetic strategy to stably knockdown eIF4E in SOM and PKCδ INs for a defined period. We used a knock-in mouse-based conditional expression of a synthetic micro-RNA specifically targeting *eif4e* mRNA^26^, consisting of *Eif4e-specific* shRNA embedded in the microRNA-30 backbone (shmiR) (Fig. 2a). shmiRs are driven by Pol II promoters and act as natural substrates in miRNA biogenesis pathways, leading to robust expression of mature shRNA and high knockdown efficiency^27^. The shmiR for *Eif4e* (shmiR-4E) is integrated in the 3’ UTR of GFP and is under transcriptional regulation of tet-responsive elements (TRE). In double transgenic SOM Cre::TRE GFP.shmiR-4E and PKCδ Cre:TRE GFP.shmiR-4E mice, we virally expressed the cre-dependent tet transactivator (tTA) in the CeL while placing the animals on Off-Dox diet for 14 days following viral delivery to mediate eIF4E knockdown (4Ekd) (Fig. 2b-c). This strategy resulted in significant reduction of eIF4E protein (Fig. 2d, Extended Data Fig. 3a-b) and subsequently, in a significant inhibition of *de novo* global translation in CeL INs (Fig. 2e, Extended Data Fig. 3c-d). MMP9, the protein product of an eIF4E-sensitive mRNA important for long-lasting synaptic plasticity in the central amygdala^28,29^ was also significantly reduced in SOM and PKCδ INs (Extended Data Fig. 3e-f). At the level of behavior, eIF4E knockdown in SOM INs did not affect spontaneous locomotion in the open field and elevated plus maze (Extended Data Fig. 4a-g). However, PKCδ 4Ekd mice, despite exhibiting normal open field activity (Extended Fig. 4h-k), explored the open arm of elevated plus maze significantly more than the control animals indicating reduced anxiety (Extended Data Fig. 4l-n). Anxiolysis with cap-dependent translation inhibition in PKCδ interneurons is consistent with a previous report that optogenetically silencing PKCδ neurons in CeL decreases anxiety^30^.

**Figure 3.**
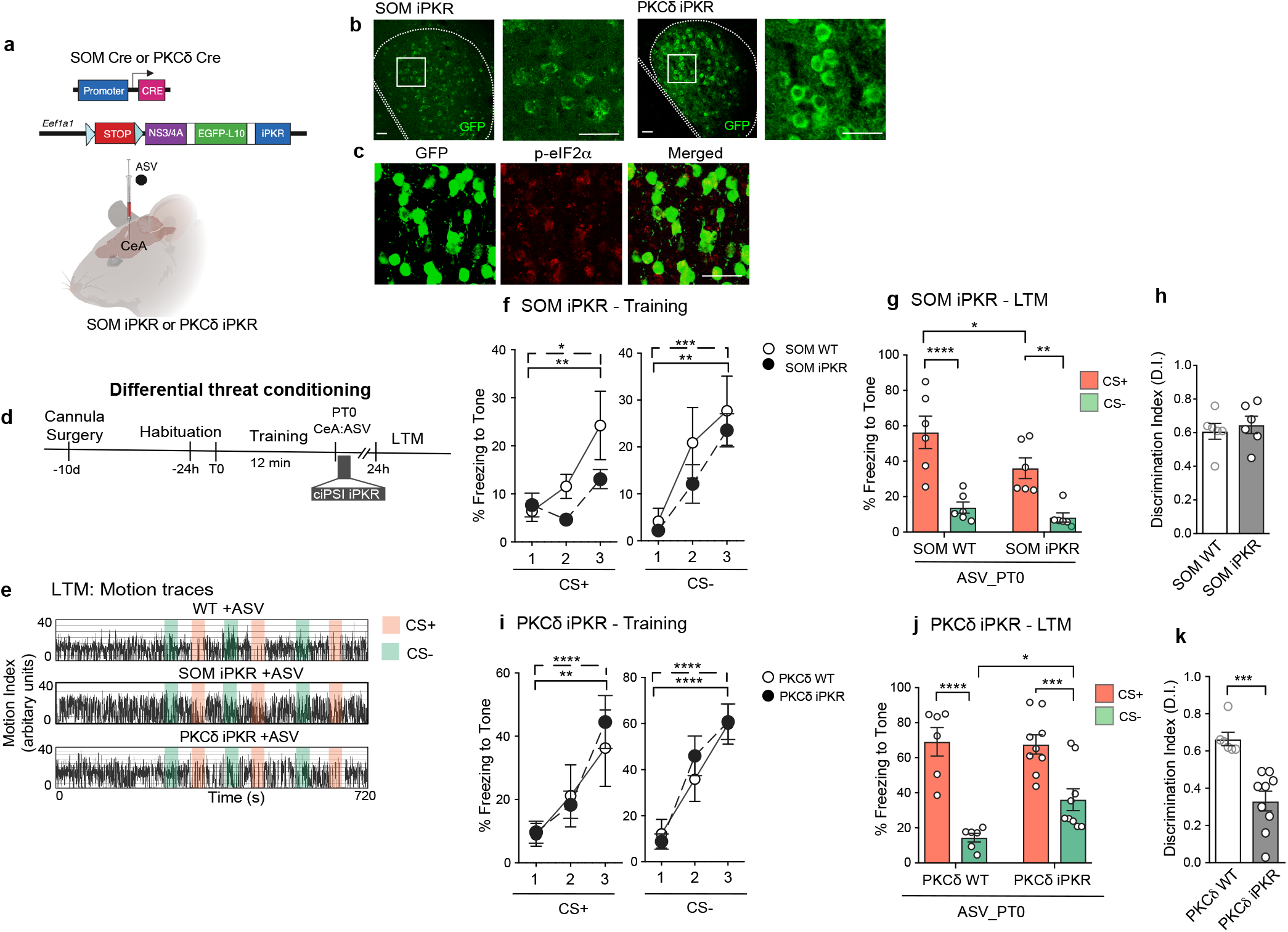
Blocking eIF2-dependent translation in specific CeL interneurons impairs consolidation of differential threat memories. a) Chemogenetic strategy for drug-inducible, cell type-specific phosphorylation of eIF2α in SOM and PKCδ neurons in CeL. b) EGFP-L10 expression in SOM iPKR and PKCδ iPKR CeL. Insets show higher magnification. c) Immunostaining for p-eIF2α showing increased eIF2α phosphorylation with drug-mediated iPKR expression. d) Behavior paradigm for differential cued threat conditioning with temporally precise protein synthesis inhibition during initial consolidation. e) Representative LTM motion traces for WT +ASV, SOM iPKR +ASV and PKCδ iPKR +ASV animals. f) Normal memory acquisition in SOM iPKR mice as they show a progressive increase in freezing response to successive presentations of the CS. CS+: F(2,28)=15.76, p<0.0001; CS-: F(2,28)=21.57, p<0.0001. n=7-9/group. g) Intra-CeL infusion of ASV decreased the threat response to CS+ in SOM iPKR animals while sparing the conditioned safety response to CS-. Genotype: F(1,20)=4.90, p=0.0376; CS: F(1,20)=36.78, p<0.0001. n=7-10/group. h) Normal discrimination index for cued threat in SOM WT and SOM iPKR animals. i) XY plot for training. Normal memory acquisition in PKCδ WT and PKCδ iPKR mice as they show progressive increase in freezing response to successive presentations of CS’s. CS+: F(2,26)=19.49, p<0.0001; CS-: F(2,26)=36.46, p<0.0001. n=6-9/group. j) Intra-CeL infusion of ASV in PKCδ iPKR mice did not affect the threat response to CS+ but significantly impaired the safety response to CS-. CS: F(1,26)=48.85, p<0.0001. n=6-9/group. k) Discrimination index for cued threat was significantly impaired in PKCδ iPKR +ASV animals compared to controls. p=0.0005. n=6-9/group. Statistical tests: RM Two-way ANOVA with Bonferroni’s post-hoc test (f, i), Two-way ANOVA with Bonferroni’s post-hoc test (g, j), Unpaired t-test (h,k). Data are presented as mean+SEM. *p<0.05, **p<0.01, ***p<0.001, ****p<0.0001. n.s. nonsignificant. Scale bar, 50 μm.

**Figure 4.**
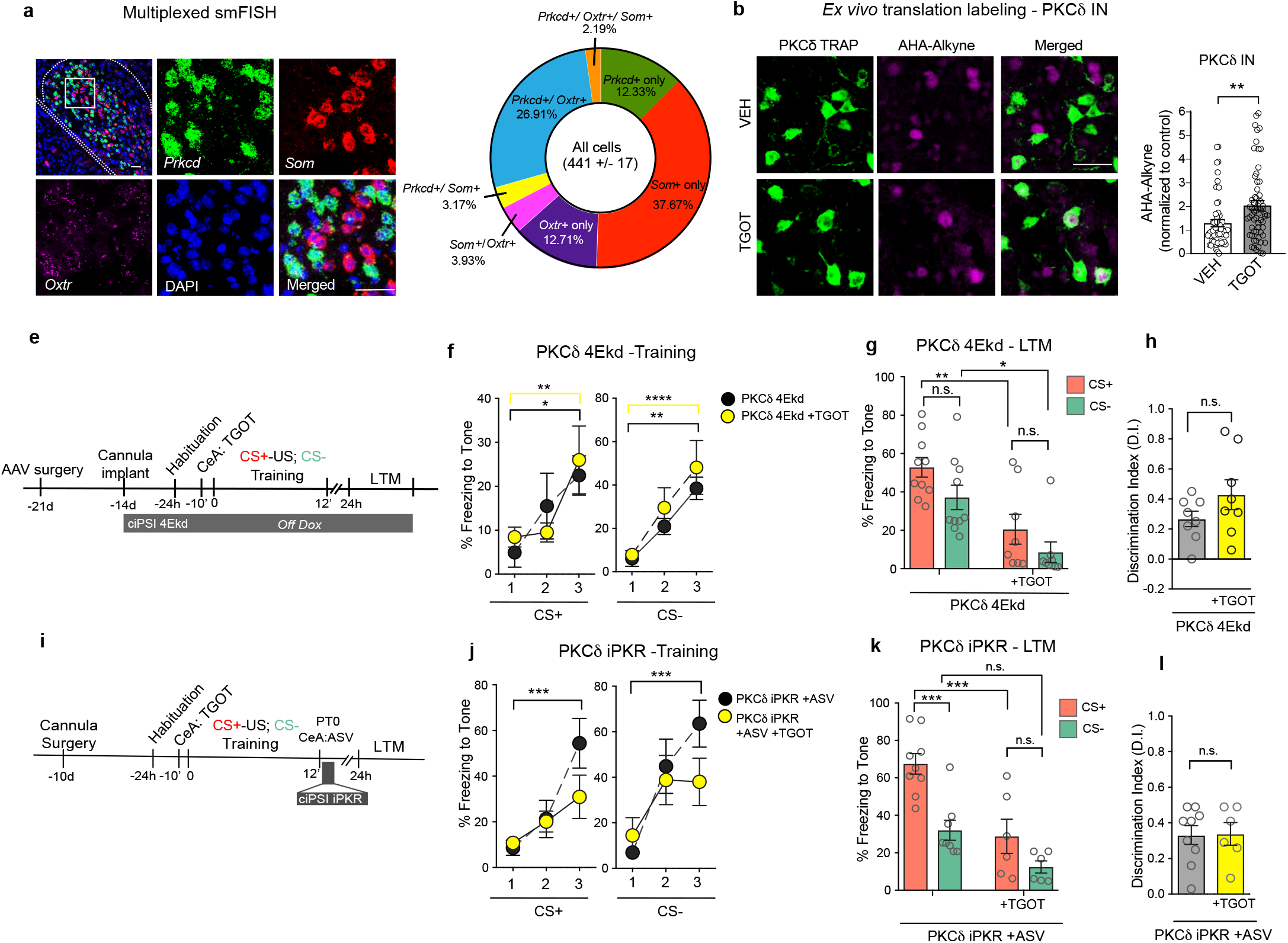
Oxytocin signaling promotes *de novo* translation in PKCδ INs and lowers both conditioned threat and safety responses. a) Multiplexed small molecule fluorescent in situ hybridization (smFISH) for *Prkcd, Som* and *Oxtr* along with DAPI nuclear counterstain. b) *De novo* translation labeling using FUNCAT in PKCδ TRAP amygdala sections. Bath application of TGOT significantly enhances *de novo* translation in PKCδ neurons. p=0.0037. n=54-61/group. e) Behavior protocol for TGOT infusion in PKCδ 4Ekd and WT animals. f) XY plot for training. Pre-training TGOT infusion in the CeL did not alter learning in PKCδ 4Ekd animals. CS+: F(2,24)=10.73, p=0.0005; CS-: F(2,24)=22.03, p<0.0001. n=6-8/group. g) Bilateral intra-CeL infusion of TGOT leads to a rescue of the conditioned safety response to CS-, but impairs the threat response to CS+. Drug: F(1,32)=23.93, p<0.0001. n=8-10/group. h) Discrimination index for cued threat was unaltered for PKCδ 4Ekd mice with TGOT infusion. i) Experimental strategy for oxytocinergic neuromodulation of PKCδ INs with impaired general translation. j) Pre-training infusion of TGOT and ASV in CeL of PKCδ iPKR mice modestly affected the learning curve for CS+ during training (left). CS+: F(2,22)=13.77, p=0.0001; CS-: F(2,22)=15.47, p=0.0001. n=6-7/group. k) Intra-CeL infusion of TGOT in translation-impaired PKCδ iPKR mice did not rescue conditioned safety response to CS- but still significantly impaired threat LTM to CS+. Genotype: F(1,25)=22.68, p<0.0001. CS: F(1,25)=17.93, p=0.0003. n=6-9/group. l) Discrimination index for cued threat was unaltered for PKCδ iPKR +ASV mice with TGOT infusion. Statistical tests: Unpaired t-test (b, h, l), RM Two-way ANOVA (f, j), Two-way ANOVA (g, k). Data are presented as mean +SEM. *p<0.05, **p<0.01, ***p<0.001, ****p<0.0001. n.s. nonsignificant. Scale bar, 50 μm.

To test whether inhibition of cap-dependent translation in CeL interneuron subtypes has any impact on long-term threat memories, we trained SOM and PKCδ animals in simple cued threat conditioning paradigm where a tone unambiguously co-terminated with a footshock (Fig. 2f, Extended Data Fig. 5a-d). Although all mice learned the CS- US association equivalently (Fig. 2g), only SOM 4Ekd mice displayed a significant LTM deficit (Fig. 2h). SOM 4Ekd mice that were placed on Dox diet for 14 days allowing eIF4E re-expression and then re-trained in the same protocol displayed complete rescue of LTM (Fig. 2i). Next, we tested 4Ekd animals in the differential threat conditioning paradigm (Fig. 2j). SOM 4Ekd mice learned equivalently to SOM WT mice (Fig. 2k). During LTM, SOM 4Ekd mice displayed a selective impairment in the conditioned threat response to CS+, yet exhibited a normal safety response to CS- and a normal cue discrimination index (Extended Data Fig. 5e, Fig. 2l-n). PKCδ 4Ekd mice also acquired differential threat associative memory normally (Fig. 2o). However, PKCδ 4Ekd mice displayed a selective impairment in the conditioned safety response to CS- despite exhibiting a normal conditioned threat response to CS+ (Fig. 2n, p), which led to a sub-optimal cue discrimination index for PKCδ 4Ekd animals (Fig. 2q). Both SOM and PKCδ 4Ekd animals displayed negligible baseline freezing during pre-CS (Extended Data Fig. 5e-h). Overall, these results show that blocking cap-dependent translation in SOM and PKCδ INs results in selective impairment in retrieval of conditioned threat and safety responses, respectively.

**Figure 5.**
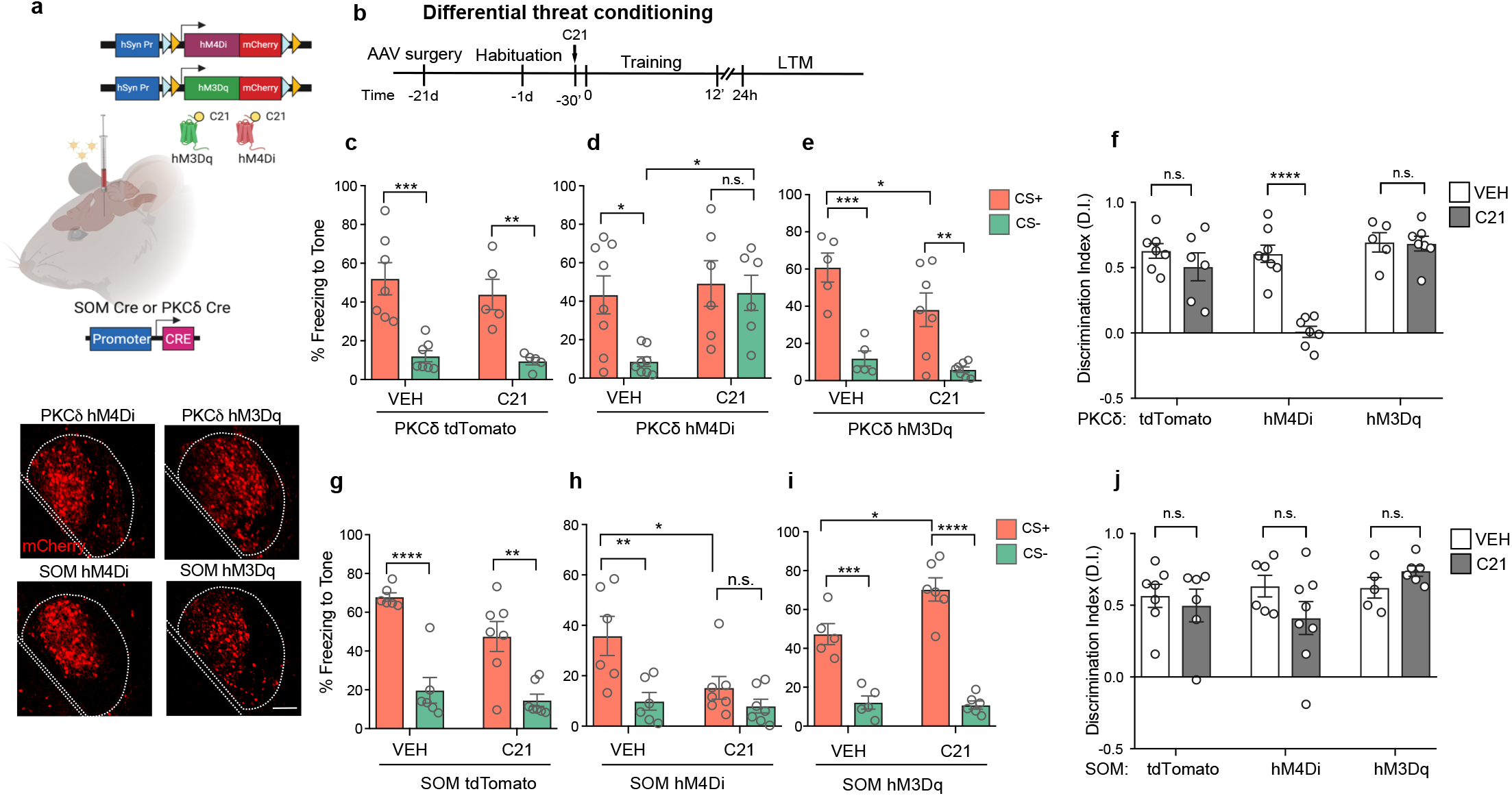
Conserved G-protein signaling pathways in CeL interneurons modulate threat and safety responses. a) Chemogenetic strategy for expressing designer G_αi_ (hM4Di) or G_αq_ (hM3Dq) protein coupled DREADD receptors in CeL PKCδ and SOM INs (top). Representative immunohistochemical images for mCherry fused to DREADD receptors in CeL PKCδ and SOM interneurons (bottom). b) Behavior paradigm design for differential threat conditioning in mice expressing DREADDs in CeL interneurons. c) Normal conditioned threat and safety responses to CS+ and CS- in PKCδ tdTomato animals injected with either vehicle (VEH) or C21. CS: F(1,20)=35.54, p<0.0001. n=5-7/group. d) The conditioned safety response to CS- is significantly impaired in PKCδ hM4Di animals administered with C21 compared to VEH control. Transgene: F(1,24)=5.702, p=0.0252; CS: F(1,24)=5.119, p=0.0330. e) The conditioned threat response to CS+ is significantly reduced in PKCδ hM3Dq animals administered with C21 compared to VEH control. Genotype: F(1,20)=4.77, p=0.041; CS: F(1,20)=38.02, p<0.0001. n=5-7/group. f) Discrimination index for cued threat is significantly impaired for PKCδ hM4Di mice +C21 mice compared to controls but unaltered for other groups. Interaction (Transgene X CS): F(2,34)=11.04, p=0.0002; Transgene: F(2,34)=16.99, p<0.0001; CS: F(1,34)=19.08, p=0.0001. n=6/group. g) Normal conditioned threat and safety responses to CS+ and CS- in SOM tdTomato animals injected with either vehicle (VEH) or C21. h) The conditioned threat response to CS+ is significantly impaired for SOM hM4Di +C21 group compared to VEH control. Transgene: F(1,22)=5.39, p=0.0299; CS: F(1,22)=11.76, p=0.0024. n=?? i) The conditioned threat response to CS+ is significantly increased in SOM hM3Dq +C21 group compared with VEH control. Interaction (Transgene X CS): F(1,18)=7.17, p=0.0153; Transgene: F(1,18)=5.70, p=0.0281. n=??/group. f) Discrimination index for cued threat is normal across all SOM groups. Two-way ANOVA with Bonferroni’s post-hoc test (c-i). Data are presented as mean +SEM. *p<0.05, **p<0.01, ***p<0.001, ****p<0.0001. n.s. nonsignificant. Scale bar, 50 μm.

To understand the contribution of time-limited *de novo* protein synthesis during the initial consolidation window following learning, we applied a knock-in mouse-based chemogenetic strategy that we recently developed^6^ to express cre-dependent and drug-inducible double-stranded RNA activated protein kinase (iPKR) in SOM and PKCδ INs^6^ (Fig. 3a). Because the iPKR mouse line also enables cre-dependent expression of EGFP-tagged ribosomal subunit L10, we detected soma-localized GFP in the CeL SOM and PKCδ neurons of SOM iPKR and PKCδ iPKR mice, respectively (Fig. 3b). *In vivo* infusion of ASV, the drug inducer of iPKR, significantly elevated phosphorylation of eIF2α (S51) in SOM and PKCδ interneurons in the CeL (Fig. 3c, Extended Data Fig. 6a-b). We then exposed the animals to the differential threat conditioning paradigm as described above, but restricted cell type-specific protein synthesis inhibition (ciPSI) to the initial consolidation period with intra-CeL infusion of ASV immediately after training (Fig. 3d). Although all mice learned equivalently during training (Fig. 3f and Fig. 3i), we found that the memory deficits were remarkably divergent in the SOM iPKR and PKCδ iPKR mice. Similar to the 4Ekd approach, we found that blocking general translation with increased eIF2α phosphorylation in SOM INs impaired the freezing response to CS+ while keeping the safety response and cue discrimination intact (Fig. 3e, g, h). On the other hand, blocking general translation with increased eIF2α phosphorylation in PKCδ INs resulted in an impaired safety response and cue discrimination, with no reduction in freezing response to CS+ during LTM (Fig. 3j-k). These findings demonstrate that the simultaneous consolidation of long-lasting threat and safety responses requires *de novo* protein synthesis in distinct populations of interneurons in centrolateral amygdala.

Oxytocinergic neuromodulation in central amygdala has been shown to be important for social discrimination as well as for suppression of conditioned freezing response^31, 32^. To determine if oxytocin acts through CeL SOM and/or PKCδ INs, we carried out multiplexed small-molecule fluorescent in situ hybridization (smFISH) in the CeL of wild-type mice using probes for the Oxytocin receptor (*Oxtr*), SOM (*Som*), and PKCδ (*Prkcd*). Of all the cells in the CeL, we found a substantial overlap between *Oxtr* and *Prkcd*, but minimal overlap between *Som* and *Oxtr* (Fig. 4a, Extended Data Fig. 7a). Approximately 27% of CeL cells were double positive for *Oxtr* and *Prkcd*, whereas 4% of all cells were *Som+/Oxtr+* and 2% cells were positive for all three markers (*Oxtr, Prkcd* and *Som*). These data show that PKCδ INs in CeL express Oxytocin receptors and are thus likely to be modulated by Oxytocin. To assess whether oxytocinergic neuromodulation increases *de novo* protein synthesis in PKCδ INs, we utilized fluorescent non-canonical amino acid tagging (FUNCAT) with the synthetic methionine analog azidohomoalanine (AHA), which gets incorporated in place of methionine in the growing polypeptide chain. Bath application of the highly selective Oxytocin receptor agonist (Thr^4^, Gly^7^)-Oxytocin (TGOT)^33^ to amygdala slices significantly increased *de novo* translation in CeL PKCδ neurons compared to vehicle treatment (Fig. 4b). These data indicate that Oxytocinergic signaling acts on CeL PKCδ INs to stimulate *de novo* protein synthesis.

Although the membrane-bound Oxytocin receptor is upstream of the protein synthesis machinery, we hypothesized that oxytocinergic stimulation might mobilize the translation apparatus and ameliorate the amnesic effects of blocking the eIF4E- and eIF2-dependent arms of translation initiation in PKCδ INs. We bilaterally infused TGOT into the CeL of PKCδ 4Ekd mice before training and assessed the effects of TGOT on differential threat memory (Fig. 4e). TGOT led to a modest increase in immobility during pre-CS of training phase (Extended Data Fig. 7b), but did not affect associative learning in PKCδ 4Ekd animals (Fig. 4f). During memory retrieval, TGOT treated mice recovered conditioned safety response to CS- but at the same time displayed reduced freezing response to CS+ that resulted in a lack of significant improvement in cue discrimination (Fig. 4g-h). The depressive effect on the threat response to CS+ is likely due to the supraphysiological effect of TGOT in CeL PKCδ INs and OxtR-expressing non-PKCδ INs, leading to disynaptic inhibition of SOM INs. We next sought to determine whether TGOT could rescue the conditioned safety response when *de novo* general translation is acutely blocked in PKCδ INs during initial consolidation using the iPKR approach (Fig. 4i). Pre-training infusion of TGOT in the CeL of PKCδ iPKR mice caused a modest increase in pre-CS freezing behavior (Extended Data Fig. 7d) and a trend-level decrease in associative learning (Fig. 4j). Both groups of mice, PKCδ iPKR and PKCδ iPKR +TGOT, were infused with ASV to block general translation immediately following training. During the LTM test, TGOT-treatment did not recover the CS- safety response; however, it significantly decreased threat response to CS+ in PKCδ iPKR animals (Extended Data Fig. 7e, Fig. 4k). As a result, the cue discrimination index did not improve for TGOT treatment (Fig. 4l). These data indicate that inhibiting general translation in PKCδ INs occludes recovery of the conditioned safety response by TGOT, and that in addition to PKCδ INs, non-PKCδ INs harboring OxtRs are likely engaged in suppressing the threat response to CS+. We next used SOM 4Ekd mice to determine whether TGOT mediates threat response inhibition by indirectly acting on SOM INs. Intra-CeL infusion of TGOT in SOM 4Ekd animals led to an increase in baseline freezing during pre-CS of training phase (Extended Data Fig. 7f) and significantly impaired CS+ threat learning despite intact nociceptive response to US (Extended Data Fig. 7g). Strikingly, TGOT treatment eliminated the freezing response to CS- in SOM 4Ekd mice, resulting in near-perfect cue discrimination, but did not reduce conditioned threat response to CS+ (Extended Data Fig. 7g-h). These findings indicate that the conditioned safety response is mediated by oxytocinergic stimulation of PKCδ INs and that the conditioned threat response to CS+ is stored in SOM INs by activating the translation machinery.

Despite the clear effects of TGOT on *de novo* translation in PKCδ INs and on conditioned safety response, it is still possible that the small fraction of cells in CeL that are non-PKCδ+ but express the Oxytocin receptor could mediate the behavioral effects of TGOT on threat and safety learning^34^. Hence, we examined the conserved cell-autonomous G_αi_ and G_αq_ protein signaling pathways in CeL interneurons using viral expression of designer receptors activated by designer drugs (DREADDs) that are based on mutant muscarinic acetylcholine receptors and couple to G proteins^35^ similar to OxtRs (Fig. 5a-b). Pretraining injection of DREADD actuator C21 did not alter associative learning in PKCδ hM4Di or PKCδ hM3Dq animals (Extended Data Fig. 8a-c). During memory retrieval, C21-treated PKCδ tdTomato control mice exhibited normal threat and safety responses to CS+ and CS- respectively (Fig. 5c), but the drug had diverse effects in PKCδ hM4Di and PKCδ hM3Dq animals. Activation of G_αi_ protein signaling pathway in PKCδ INs in PKCδ hM4Di +C21 mice led to a selective impairment in safety response to CS- and cue discrimination (Fig. 5d, f) whereas activating the G_αq_ pathway in PKCδ INs reduced the threat response to CS+ (Fig. 5e-f). C21 had no effect on baseline freezing during pre-CS of both memory acquisition and retrieval phases (Extended Data Fig. 8 d-e). These data indicate that G_αi_ protein signaling that generally leads to inhibition of adenylyl cyclase mirrors the effect of blocking *de novo* protein synthesis in PKCδ INs and that G_αq_ pathway activation in PKCδ INs is consistent with the effects of intra-CeL TGOT infusion, suggesting G_αq_ coupling of OxtRs in these CeL INs.

Finally, we examined the effects of activating G_αi_ and G_αq_ signaling pathways in CeL SOM INs. C21 treatment did not alter memory acquisition in SOM tdTomato, SOM hM4Di and SOM hM3Dq animals (Extended Data Fig. 8 f-h). During memory retrieval, C21 had no effect on threat and safety responses in control SOM tdTomato animals (Fig. 5g), but the drug exerted opposite effects on SOM hM4Di and SOM hM3Dq animals. We found that C21 treatment significantly decreased the conditioned threat response to

CS+ in SOM hM4Di mice (Fig. 5h). This threat response deficit caused by activating G_αi_ protein signaling in SOM INs is consistent with behavioral effects of *de novo* translation inhibition in these CeL INs. In contrast, increasing neuronal activity in SOM INs by activating G_αq_ pathway resulted in enhanced CS+ LTM (Fig. 5i) supporting bidirectional modulation of the threat response with chemogenetic manipulation of conserved G-protein signaling in SOM INs, consistent with previous findings^1,3^. C21 did not alter baseline freezing during pre-CS of both memory acquisition and retrieval phases (Extended Data Fig. 8 i-j).

Past studies have reported enhancement of long-term spatial and threat memories by relieving translation repression with constitutive deletion of genes encoding eIF2α kinases such as GCN2 and PKR^36,37^ or by administering ISRIB, an eIF2B activator^38^. Likewise, constitutive deletion of the gene encoding the eIF4E repressor 4E-BP2 results in enhanced conditioned taste aversion memory^39^ whereas acute intra-amygdalar infusion of 4EGI-1, an inhibitor of eIF4E-eIF4G interaction, blocks threat memory consolidation^40^. In both simple and differential threat conditioning paradigms, our results show that eIF2-as well as eIF4E-dependent translation programs in CeL SOM INs are required for the conditioned threat response, which indicates that SOM INs are the primary CeL locus for storage of cued threat memory. Our findings are consistent with studies showing long-lasting synaptic potentiation in CeL SOM interneurons following threat learning that lasts at least 24 hours^13^. Moreover, expressing biallelic phosphomutant eIF2α in SOM INs brainwide results in enhanced cued and contextual LTM^41^. In a contrasting but complementary role, *de* novo translation in PKCδ INs serves to store the conditioned safety response. Our findings thus support a working model where CeL SOM and PKCδ INs simultaneously store threat and safety cue associated memories by changing the cellular translation landscape (Extended Data Fig. 9).

Threat generalization due to impaired safety response is a hallmark feature of PTSD^**6**^. How does threat generalization between similar stimuli occur? During auditory cued threat conditioning, information about the CS arrives in LA from auditory cortex and auditory thalamus. Cells in the auditory cortex undergo dynamic receptive field plasticity characterized by a shift in their tuning to the CS+ frequency as the new best frequency (BF)^42^. The medial part of the auditory thalamus, i.e. medial geniculate (MGm), on the other hand is broadly tuned to multiple tone frequencies and increasing CREB expression in this region results in broadening of tone threat generalization^43^. Overtraining or increasing US intensity has also been shown to increase auditory threat generalization^44^. In particular, cells in the LA shift the threat response from cue-specific to cue generalization depending on the US intensity^45^. CeL PKCδ INs express calcitonin gene-related peptide (CGRP) receptors and are direct recipients of US related nociceptive input from the parabrachial nucleus^46^. CeL PKCδ INs in turn convey the processed US related information to LA and are instructive for associative threat learning^4^. Our current findings that blocking neuronal activity and *de novo* protein synthesis in CeL PKCδ INs disrupts the acquisition and consolidation of long-term conditioned response inhibition to CS- is in agreement with the US processing feature of these types of neurons. It is possible that during differential threat learning, nociceptive afferents from PBN activate PKCδ INs to cues of similar frequency, but oxytocinergic neuromodulation of PKCδ INs actively triggers protein synthesis-dependent form of long-term depression (LTD) and occludes the generation of the threat response to CS-. Oxytocin-mediated LTD has been reported for the medial amygdala, a brain region proximal to CeL, during long-term social recognition^47^ suggesting that CeL PKCδ INs might express similar form of synaptic plasticity under the neuromodulatory influence of oxytocin.

To conclude, our study provides the first evidence that disruption of protein synthesis in discrete interneuron subpopulations in the centrolateral amygdala impairs associative memories related to threat and safety, which may contribute to maladaptive behavior in memory disorders such as PTSD.

## Materials and Methods

### Animals

Mice were provided with food and water ad libitum and were maintained in a 12h/12h light/dark cycle at New York University at stable temperature (78°F) and humidity (40 to 50%). All mice were backcrossed to C57Bl/6J strain for at least 5 generations. Both male and female mice, aged 3-6 months, were used in all experiments. Somatostatin IRES-cre knockin mice (SOM Cre; stock #013044) were obtained from Jackson labs. PKCδ:: GluClα-iCre BAC transgenic mice (PKCδ Cre; Haubensak et al. 2010) were generated by GENSAT and kindly provided by Dr. David Anderson (Caltech). Cre reporter lines including Floxed TRAP (stock #022367) mice expressing GFP-L10 fusion protein in a cre-dependent manner, and Floxed tdTomato mice (Ai14; stock #007908) that express tdTomato in a cre-dependent manner were obtained from Jackson labs. Col1a1^TRE GFP.shmiR-4E.389^ mice were generated as previously described (Cell Reports 2012). Floxed iPKR (Eef1a1^LSL.NS3/4.TRAP.iPKR^) mice were generated as previously described (Shrestha 2020). SOM Cre and PKCδ Cre mice were crossed with floxed Col1a1^TRE GFP.shmiR-4E^ mice to generate transheterozygote SOM.TRE GFP^shmiR-eIF4E^ and PKCδ.TRE GFP^shmiR-eIF4E^ mice respectively. Likewise, SOM Cre and PKCδ Cre mice were crossed with floxed iPKR mice to generate transheterozygote SOM iPKR and PKCδ iPKR mice respectively. SOM tdTomato and PKC5 tdTomato mice were generated by crossing SOM Cre and PKCδ Cre with floxed tdTomato reporter line, whereas PKCδ TRAP mice were generated by crossing PKCδ Cre line with floxed TRAP mice. SOM.tdTomato/TRE GFP^shmiR-eIF4E^ and PKCδ. tdTomato/TRE GFP^shmiR-eIF4E^ mice were generated by breeding SOM.TRE GFP^shmiR-eIF4E^ and PKCδ.TRE GFP^shmiR-eIF4E^ mice with homozygous floxed tdTomato reporter line. Wildtype C57Bl/6J mice (stock #000664) were purchased from Jackson labs.

### Drugs and chemicals

Doxycycline was added to rodent chow at 40 mg/kg (Bio-Serv, F4159). This doxycycline diet was provided to SOM 4Ekd, PKCδ 4Ekd, and control SOM vGFP and PKCδ vGFP mice starting from the day of surgery for 7d and to SOM 4Ekd re-training group for 14d after LTM1 ad libitum. Asunaprevir (ASV, ChemExpress) was dissolved in DMSO to a stock concentration of 10 mM and diluted in sterile saline to 100 nM. 0.5 μl of this drug was intracranially infused into the central amygdala (−1.22 mm anterioposterior AP, +/-3.00 mm ML, −4.60 mm DV) of SOM.iPKR and PKCδ.iPKR animals using an injection cannula inserted into the stainless steel guide cannula (Plastics One). ASV infusion was carried out at 0.125 μl/min using an injection cannula extending out of PE50 tubing attached to a 5 μl Hamilton syringe (Hamilton) using a PHD 2000 Infusion Pump (Harvard Apparatus). After injection, the injection cannula was kept in place for 1 min before its withdrawal. Puromycin (Sigma, P8833) was dissolved in ddH_2_O at 25 μg/μl, and this stock was freshly diluted in saline to 10 μg/μl for SUnSET assays *in vivo*. Thr^4^, Gly^7^-Oxytocin (TGOT) (VWR, # H-7710.0005BA) was dissolved in ddH_2_O at a concentration of 0.4 mM for stock solution and was further diluted in saline to a final concentration of 16 μM for intra-CeL injections (7 ng in 0.5 μl). For bath application in slices, 0.4 μM TGOT was added to ACSF. OTA (L-368,899) (Tocris, #2641) was dissolved in ddH_2_O at 10 mg/ml and was further diluted to 2 mg/ml in saline for intra-CeL injections (1 μg in 0.5 μl). Azidohomoalanine (AHA) (Fisher, # NC0667352) was dissolved in ddH_2_O at a stock concentration of 100 mM and diluted to 1 mM in ACSF. Digitonin (Sigma, D141) was dissolved in ddH_2_O at 5% w/v to prepare the stock solution, which was diluted to 0.0015% w/v in 0.1M PBS. Stock solution of aqueous 32% paraformaldehyde (EMS, 15714) was freshly diluted to 4% in 01.M PBS for transcardial perfusions and post-fixation of brain slices. The DREADD actuator, agonist C21 (Tocris), was dissolved in DMSO at 40 mg ml-1 concentration, freshly diluted in saline and administered to mice at 1 mg kg-1 intraperitoneally.

### Stereotaxic surgeries

Mice were anesthetized with the mixture of ketamine (100 mg/kg) and xylazine (10 mg/kg) in sterile saline (i.p. injection). Stereotaxic surgeries were carried out on the Kopf stereotaxic instrument (Model 942), which was equipped with a microinjection unit (Model 5000). Viral vectors were injected intracranially using 2.0 μl Neuros syringe (Hamilton, #65459-02). Postoperative analgesia was delivered using subcutaneous injections of ketoprofen (3 mg/kg) for 3 days starting from the day of surgery. To generate SOM.4Ekd and PKCδ.4Ekd mice, 300 nl of AAV9.CAG Pr.DIO.tTA (1.0 × 10^13 GC/ml; Vigene) was injected into the centrolateral amygdala (CeL) [-1.22 mm anterior posterior (AP), +/- 3.00 mm mediolateral (ML) and −4.60 mm dorsoventral (DV)] of double transheterozygote SOM.TRE^GFP.shmiR-4E^ or PKC5. TRE ^GFP.shmιR-4E^ mice. The plasmid encoding tet transactivator in a cre-selective manner and under the transcriptional control of CAG promoter (pAAV.CAG Pr.DIO.tTA) was kindly provided by Hongkui Zeng (Allen Institute for Brain Science). For DREADD experiments, SOM and PKCδ Cre mice were injected with 300 nl of AAV8.hSyn Pr.DIO.hM3Dq-mCherry (≥ 4×1O^12^ vg/mL; Addgene #44361-AAV8) or AAV9.hSyn Pr.DIO.hM4Di-mCherry (≥ 1×10^13^ vg/mL, Addgene # 44362-AAV9). For controls, wild-type SOM and PKCδ mice were injected in CeL with 100 nl of AAV.CAG Pr.DIO.GFP (3.33 × 10^13 GC/ml) to generate SOM vGFP and PKCδ vGFP mice. Behavior and histology experiments for all viral vector injected animals were carried out 2-3 weeks after surgery. A cohort of SOM.iPKR and PKCδ.iPKR mice were injected with bilaterally in CeL with 200 nl of AAV.Eef1a1 Pr.DIO.EGFPL10a (7 × 10^^12 GC/ml, Addgene) for immunohistochemistry experiment;. Intracranial cannula implant surgeries were carried out using custom-designed guide cannulas (Plastics One) along with a skull screw (1.6 mm shaft) to stabilize the dental cement, Metabond quick adhesive cement (Parkell S380) encapsulating the skull surface. For *in vivo* surface labeling of translation (SUnSET), SOM.4Ekd, PKCδ.4Ekd and control animals were implanted with a 23 gauge stainless steel guide cannula in right CeL (−1.22 mm AP, +3.00 mm ML and −2.40 mm DV) for puromycin infusion using an internal cannula with 2 mm projection. For TGOT experiments, SOM.4Ekd and PKCδ.4Ekd were bilaterally implanted with the guide cannulas in CeL (−1.22 mm AP, +/-3.00 mm ML and −2.40 mm DV). For OTA infusion experiments, wildtype C57Bl/6J mice were bilaterally implanted with guide cannulas in CeL. Similarly, SOM.iPKR and PKCδ.iPKR mice were also implanted with the 23 gauge stainless steel cannulas in CeL bilaterally for ASV infusions.

### Behavior

All behavior sessions were conducted during the light cycle. Both male and female mice were included in all behavior experiments. SOM.4Ekd, PKCδ.4Ekd and control mice were trained in threat conditioning paradigms after 14 days of eIF4E knockdown (Off Dox). A separate group of SOM.4Ekd, PKCδ.4Ekd and control mice were tested in the open field arena and elevated plus maze test after the same duration of eIF4E knockdown. SOM.iPKR and PKCδ.iPKR animals were trained in threat conditioning paradigms 10 days after cannula implant surgeries to allow time for recovery.

### Open field activity

Mice were placed in the center of an open field (27.31 × 27.31 × 20.32 cm) for 15 min during which a computer-operated optical system (Activity monitor software, Med Associates) monitored the spontaneous movement of the mice as they explored the arena. The parameters tested were: distance traveled, and the ratio of center to total time.

### Elevated plus maze

The plus maze consisted of two open arms (30 cm × 5 cm) and two enclosed arms of the same size with 14-cm high sidewalls and an endwall. The arms extended from a common central square (5 cm^2^ × 5 cm^2^) perpendicular to each other, making the shape of a plus sign. The entire plus-maze apparatus was elevated to a height of 38.5 cm. Testing began by placing a mouse on the central platform of the maze facing the open arm. Standard 5-min test duration was applied and the maze was wiped with 30% ethanol in between trials. Ethovision XT13 software (Noldus) was used to record the time spent on open arms and closed arms, total distance moved, and number of open arm and closed arm entries.

### Simple cued threat conditioning

Mice were habituated for 15 min in the threat conditioning chambers housed inside sound attenuated cubicles (Coulbourn instruments) for 1 day. The habituation and training context included a metal grid floor and a white houselight. For simple threat conditioning, mice were placed in the context for 270s and then presented twice with a 5kHz, 85 dB pure tone for 30s that co-terminated with a 2s 0.5mA footshock. The intertrial interval (ITI) was 2 min and after the second tone-shock presentation, mice remained in the chamber for an additional 120s. Cued threat conditioning (cTC) LTM was tested 24h after training, in a novel context (Context B: vanilla scented cellulose bedding, plexiglas platform, and red houselight) with three presentations of paired tone (conditioned stimulus, CS). Freezing behavior was automatically measured by Freeze Frame software (ActiMetrics) and manually re-scored and verified by an experimenter blind to the genotype/drug. Motion traces were generated using the Freeze Frame software.

### Differential cued threat conditioning

For standard differential threat conditioning, mice were placed in the training context for 250s and then trained with interleaved presentations of three paired tones or CS+ (7.5 kHz pulsatile tone, 50% duty cycle) that co-terminated with a 0.5mA footshock and three unpaired tones or CS- (3 kHz pure tone) in the training context with variable ITI. Specifically, the CS+ (7.5 kHz) was presented at 270, 440 and 570s and were paired with a footshock, whereas the 3 kHz pure tone occurred at 370, 520 and 660s. The following day, cued threat discrimination (cTD) LTM was tested with 3 interleaved presentations of CS+ and CS- tones with the order reversed from the training day and with variable intertrial intervals. Specifically, the 3kHz CS- tone was presented at 250, 380 and 550s, whereas the 7.5kHz pulsed tone was presented at 310, 450 and 630 s. All tones lasted for 30s. After the last CS- tone, mice remained in the testing context for an additional 60s. When specifically stated, the CS- tones were assigned as 1 kHz pure tone. Box-Only control group were placed in the training context for the same duration as the cTD (Paired) group but they did not receive any footshock or were exposed to either CS+ or CS-. Unpaired control group were presented with three interleaved presentations of CS+ and CS- like the cTD (Paired) group, however the US was presented in between the CSs with no tone-shock contingency. All groups of mice (Box-Only, Paired and Unpaired) were tested the following day with three presentations of CS+ and CS- in reverse sequence compared to the training day. Freezing behavior was automatically measured by Freeze Frame software (ActiMetrics) and manually re-scored and verified by an experimenter blind to the genotype/drug. Motion traces were generated using the Freeze Frame software. Discrimination index was calculated as follows:

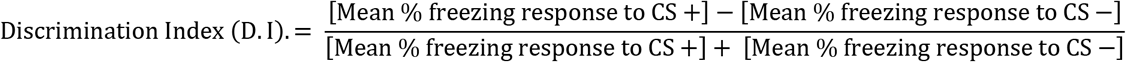

### Western immunoblot

Mice were euthanized by cervical dislocation. 300 μm-thick brain slices containing amygdala [Bregma - 1.22 mm to −2.06 mm] were prepared in cold (4°C) carbooxygenated (95% O_2_, 5% CO_2_) cutting solution ( 110 mM sucrose, 60 mM NaCl, 3 mM KCl, 1.25 mM NaH_2_PO_4_, 28 mM NaHCO_3_, 5 mM Glucose, 0.6 mM Ascorbate, 7 mM MgCl_2_ and 0.5 mM CaCl_2_) using a VT1200S vibratome (Leica). The amygdala was micro-dissected from the brain slices and sonicated in ice-cold homogenization buffer (10 mM HEPES, 150 mM NaCl, 50 mM NaF, 1 mM EDTA, 1 mM EGTA, 10 mM Na4P2O7, 1% Triton X-100, 0.1% SDS and 10% glycerol) that was freshly supplemented with 10 μl each of protease inhibitor (Sigma) and phosphatase inhibitor (Sigma) per ml of homogenization buffer. Protein concentrations were measured using BCA assay (GE Healthcare). Samples were prepared with 5X sample buffer (0.25 M Tris-HCl pH6.8, 10% SDS, 0.05% bromophenol blue, 50% glycerol and 25% - ß mercaptoethanol) and heat denatured at 95°C for 5 min. 40 μg protein per lane was run in pre-cast 4-12% Bis-Tris gels (Invitrogen) and subjected to SDS-PAGE followed by wet gel transfer to PVDF membranes. After blocking in 5% non-fat dry milk in 0.1M PBS with 0.1% Tween-20 (PBST), membranes were probed overnight at 4°C using primary antibodies (rabbit anti-p S6 (S235/236) 1:1000 (Cell Signaling #4858), rabbit anti-p-S6 (S240/244) 1:1000 (Cell Signaling #5364), mouse anti-S6 1:500 (Cell Signaling #2317), rabbit anti-p-S6K1 Thr389 1:500 (Cell Signaling #9205), rabbit anti-S6K1 1:500 (Cell Signaling #2708), rabbit anti-p eIF2α Ser51 1:300 (Cell Signaling #9721), rabbit eIF2α 1:1000 (Cell Signaling #9722), mouse anti-ß tubulin 1:5000 (Sigma #T8328) and mouse anti-ß actin 1:5000 (Sigma #A1978). After washing 3 times in 0.1% PBST, membranes were probed with horseradish peroxidase-conjugated secondary IgG (1:5000) (Millipore) for 1h at RT. Signals from membranes were detected with ECL chemiluminescence (Thermo Pierce) using Protein Simple instrument. Exposures were set to obtain signals at the linear range and then normalized by total protein and quantified via densitometry using ImageJ software.

### *In vivo* surface labeling of translation (SUnSET)

Awake behaving mice with intracranial cannula implants were infused with 5 μg puromycin (0.5 μl, 10 μg/μl) in the central amygdala using PHD2000 infusion pump and Hamilton 5.0 μl syringe. Mice were returned to the home cage and translation labeling with puromycin was carried out for 1h. Mice were deeply anesthetized with a mixture of ketamine (150 mg/kg) and xylazine (15 mg/kg), and transcardially perfused with 0.1M PBS, 0.0015% digitonin followed by 4% paraformaldehyde (PFA) in PBS. Brains were extracted and postfixed in 4% PFA for 24h, followed by immunohistochemistry.

### Fluorescent Non-canonical amino acid tagging (FUNCAT)

300 μm-thick brain slices containing amygdala [Bregma −1.22 mm to −1.70 mm] were prepared in cold (4°C) carbooxygenated (95% O_2_, 5% CO_2_) cutting solution ( 110 mM sucrose, 60 mM NaCl, 3 mM KCl, 1.25 mM NaH_2_PO_4_, 28 mM NaHCO_3_, 5 mM Glucose, 0.6 mM Ascorbate, 7 mM MgCl_2_ and 0.5 mM CaCl_2_) using a VT1200S vibratome (Leica). Slices were recovered in oxygenated ACSF solution at 32°C for 2h before 1mM AHA was added to the ACSF solution for additional 2.5 h. In the last 30 min, TGOT or vehicle was added to the buffer. Following incubation, the amygdala was dissected out and submerged in 4% paraformaldehyde at 4°C overnight. The fixed micro-slices were embedded in 3% agarose, re-cut into 40 μm sections using a vibratome, and stored at −20 °C in the cryosolution overnight. The amygdala sections were blocked and permeabilized in 5% IgG protease-free BSA, 3% normal goat serum, 0.3% Triton X-100 in 1X PBS at RT for 90 min with agitation. The brain sections were then subjected to click chemistry using Cell Reaction kit (Thermo Fisher #C10269) with 25 μM Alkyne Alexa fluor 405 (Click Chemistry tools # 1309-1), CuSO4 and additive overnight at 4°C. The following day, the sections were washed 3 times with 1X PBS for 10 min per wash. Sections were blocked with 1% NGS in 1X PBS for 1h followed by incubation with primary antibodies overnight at 4°C for standard immunohistochemistry.

### Immunohistochemistry

Mice were deeply anesthetized with a mixture of ketamine (150 mg/kg) and xylazine (15 mg/kg), and transcardially perfused with 0.1M PBS followed by 4% paraformaldehyde in PBS. Brains were removed and postfixed in 4% PFA for 24h. 40 μm free-floating coronal brain sections containing amygdala were collected using Leica vibratome (VT1000s) and stored in 1X PBS containing 0.05% Na-azide at 4°C. After blocking in 5% normal goat serum in 0.1M PBS with 0.1% Triton X-100, brain sections were probed overnight with primary antibodies (chicken anti-EGFP (abcam #ab13970 1:500; for PKCδ TRAP, SOM 4Ekd and PKCδ 4Ekd brain sections), rabbit anti-EGFP 1:300 (Thermo Fisher #G10362; for SOM iPKR and PKCδ iPKR brain sections), rabbit anti-pS6 (S235/6) 1:1000 (Cell Signaling #4858), rabbit anti-p-eIF2α S51 1:300 (Cell Signaling #9721), rabbit anti-eIF4E 1:500 (Bethyl #A301-153A), rabbit anti-Mmp9 1:300 (abcam #ab38898), mouse NeuN 1:2000 (Millipore Sigma #MAB377), chicken anti-Somatostatin 1:300 (Synaptic Systems #366 006), rabbit anti-PKCδ 1:250 (abcam #ab182126), guinea pig anti-RFP 1:500 (Synaptic systems #390 004), and mouse anti-puromycin 1:1000 (Millipore Sigma #MABE343). After washing three times in 0.1M PBS, brain sections were incubated with Alexa Fluor conjugated secondary antibodies 1:200 (Abcam #ab175674, #ab175651; Thermo Fisher #A-111034, #A11012, #A21245, #A11073, #A121236, #A21206) in blocking buffer for 1.5h at RT, and mounted using Prolong Gold antifade mountant with DAPI or without DAPI (Life Technologies #P36931, #P36930).

### Single molecule fluorescence in situ hybridization

Mouse brains were collected through flash freezing in OCT Tissue Tek medium (VWR #25608-930) in dry ice. Using a cryostat, each brain was serially sectioned at 20 μm and thaw-mounted onto Superfrost plus slides spanning AP −1.22 mm to AP −1.70 mm. Slides were stored at −80°C. Single molecule fluorescent in situ hybridization (smFISH) was performed using a RNAscope fluorescent multiplex kit (ACD Bio #320850). OxtR (#41271), SOM (#404631-C2), GFP (#409011-C3), eIF4E (#499691-C2) and Prkcd (#44191-C3) probes were purchased from the Advanced Cell Diagnostics catalog. Brain sections were fixed in 4% paraformaldehyde for 15 min and then washed in 50%, 70%, 100% and 100% ethanol for 5 min each. Slides were dried for 10 min and hydrophobic barrier drawn around the sections using ImmEdge hydrophobic barrier pen (ACD Bio #310018). Proteins were digested using protease solution (Protease IV) for 30 min at RT. Immediately afterward, slides were washed twice in 0.1M PBS. C1, C2 and C3 probes were heated in 40°C water bath for 10 min, and brought to RT for additional 10 min. Probes were applied to the slides in a humidified incubator (ACD Bio #321711) for 2h. Slides were rinsed twice in RNAscope wash buffer and then underwent the colorimetric reaction steps according to manufacturer’s instructions using AMP4-Alt C (C1, red; C2, far red; C3, green). After the final wash buffer, slides were immediately coverslipped using Prolong Gold Antifade mounting medium with DAPI.

### Image analysis

Imaging data for the whole coronal brain section were acquired using Olympus slide scanner (VS120) for qualitative visualization of transgene expression and viral gene targeting, and analyzed in ImageJ using the BIOP VSI reader plugin. Imaging data from immunohistochemistry and smFISH experiments were acquired using an SP8 confocal microscope (Leica) with 20X objective lens (with 1X or 2X zoom) and z-stacks (approximately 6 optical sections with 0.563 μm step size) for three coronal sections per mouse from AP −1.22 mm to −1.70 mm (n=3 mice) were collected. Imaging data was analyzed with ImageJ using the Bio-Formats importer plugin. Maximum projection of the z-stacks was generated followed by manual outline of individual cells and mean fluorescence intensity measurements using the drawing and measure tools. Mean fluorescence intensity values for all cell measurements were normalized to the mean fluorescence intensity for controls.

### Statistics

Statistical analyses were performed using GraphPad Prism 8 (GraphPad software) for all datasets. Data are expressed as mean +/- SEM. Data from two groups were compared using two-tailed unpaired Student’s t test. Multiple group comparisons were conducted using one-way ANOVA, or two-way ANOVA, with post hoc tests as described in the appropriate figure legend. Statistical analysis was performed with an α level of 0.05. p values <0.05 were considered significant.

---

**Extended Data Figure 1.**
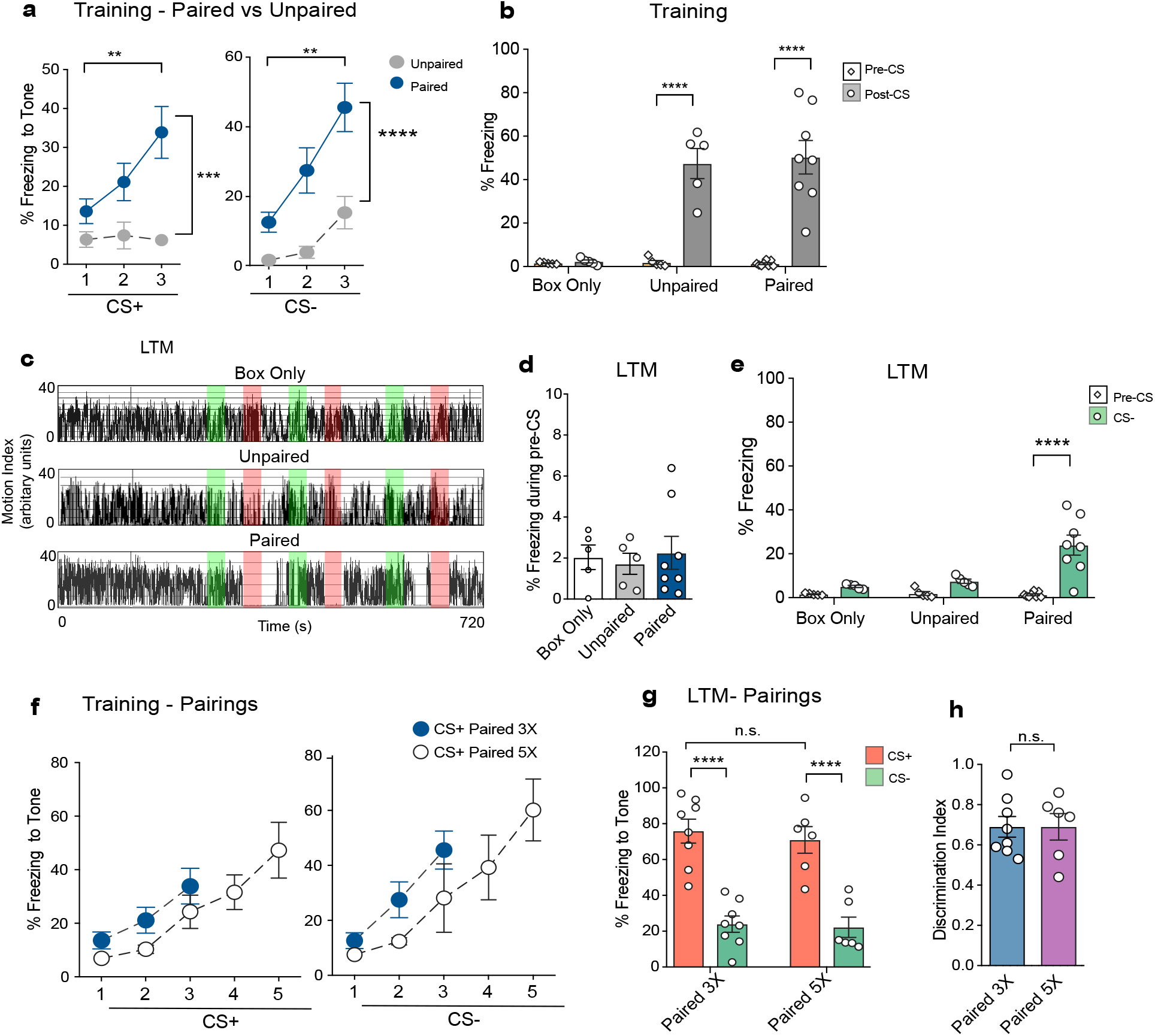
Differential cued threat conditioning. a) Paired group learned the association between CS+ and US and showed increasing freezing response to successive CS presentations whereas Unpaired group did not associate CS+ with US. RM Two-way ANOVA with Bonferroni’s post-hoc test. CS+: F(1,11)=11.40, p=0.0062; CS-: F(1,11)=11.12, p=0.0067. n=5-8/group. b) Both Paired and Unpaired groups, but not Box-Only group, increased freezing levels during post-tone period compared to pre-tone period. Two way ANOVA with Bonferroni’s post hoc test. Interaction (Training X Epoch): F(2,30)=13.94, p<0.0001, Training: F(2,30)=13.86, p<0.0001, Epoch: F(1,30)=60.38, p<0.0001. n=5-8/group. c) Representative motion traces for Box-Only, Unpaired and Paired groups during LTM. d) Freezing response during pre-CS of LTM test is low and comparable across all three groups. One-way ANOVA. n.s. e) Animals in the Paired group freeze significantly higher during CS- than during the pre-tone period. Two-way ANOVA with Bonferroni’s post-hoc test. Interaction (Training X Epoch): F(2,30)=9.12, p=0.0008, Training: F(2,30)=8.38, p=0.0013, Epoch: F(1,30)=23.97, p<0.0001. f) Increasing the number of CS- US pairs from 3 to 5 pairings during training led to a continu ed escalation of freezing response to successive presentations of CS’s. n=6-8/group. g) Paired 5X group displayed equivalent conditioned threat response and safety response to CS+ and CS- respectively as paired 3X group. Two way ANOVA with Bonferroni’s post-hoc test. Pairings: F(1,24)=0.2942, p=0.593; CS: F(1,24)=66.46, p<0.0001. n=6-8/group. h) Discrimination index for cued threat in Paired 5X group was unaltered compare to Paired 3X group. Unpaired t-test, n.s. Data are presented as mean +SEM. *p<0.05, **p<0.01, ***p<0.001, ****p<0.0001. n.s. non significant.

**Extended Data Figure 2.**
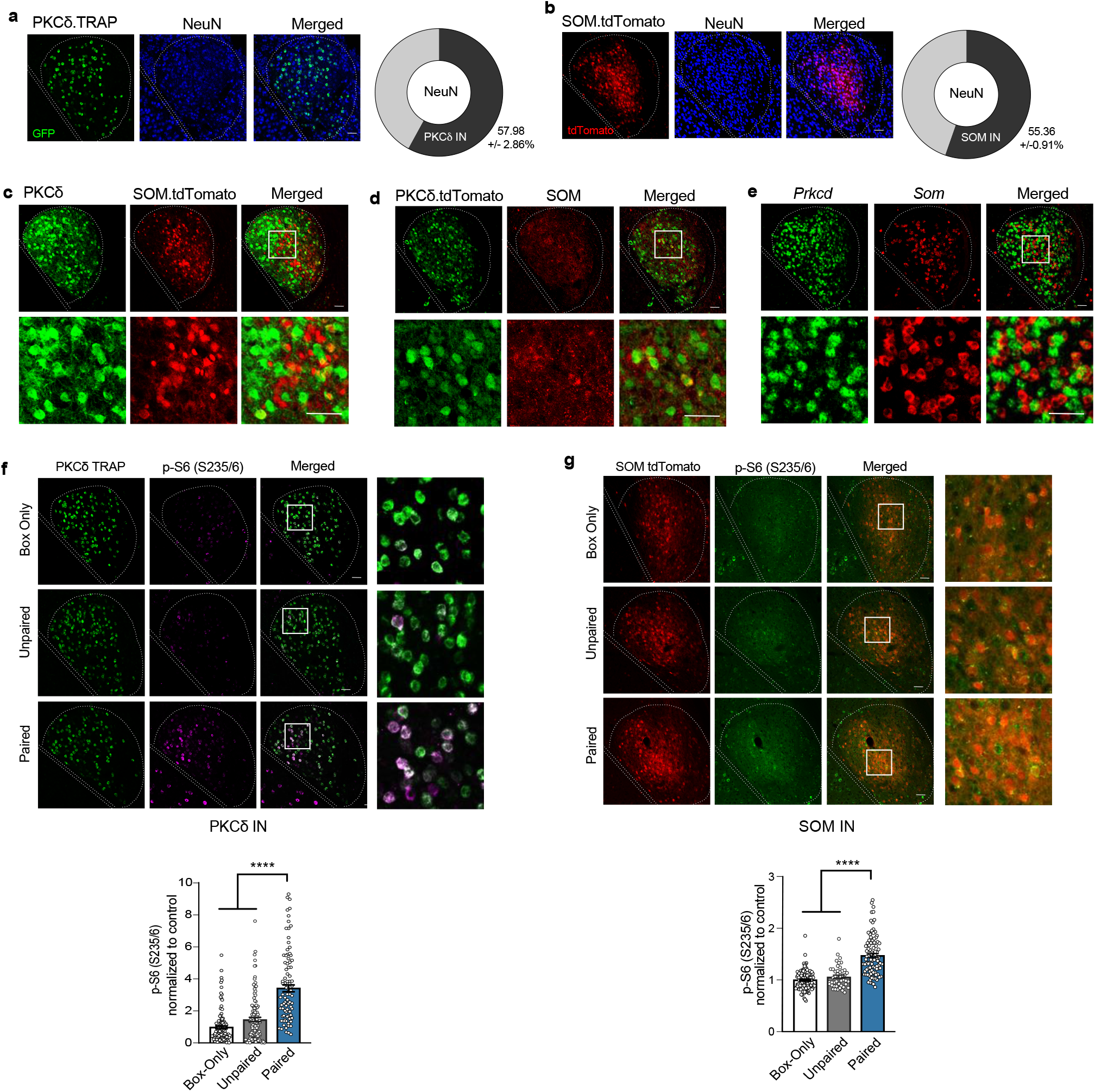
Distinct interneuron subpopulations in centrolateral amygdala. a) Co-immunostaining for GFP and neuronal marker, NeuN, in PKCδ TRAP amygdala sections. 57.96 +2.86% of all neurons in centrolateral amygdala are PKCδ positive interneurons, n=3/group. b) Co-immunostaining for tdTomato and NeuN in SOM tdTomato amygdala sections. SOM interneurons constitute 55.36 +0.91% of all neurons in CeL. n=3/group. c) Immunohistochemistry for PKCδ in SOM tdTomato brain sections shows largely non-overlapping expression of PKCδ in SOM Cre expressing cells in CeL. d) Immunohistochemistry for SOM in PKCδ tdTomato brain sections also shows largely non-overlapping populations but the subcellular distribution of SOM in neuronal processes makes it difficult to analyse the extent of SOM co-expression in PKCδ Cre expressing cell populations, e) Multiplexed smFISH for *Prkcd* and *Som* showing mutually exclusive interneurons in CeL expressing these two mRNA populations, f) Immunohistochemistry data for PKCδ TRAP amygdala sections showing expression of p-S6 (S235/6) in PKCδ neurons in CeL across three groups (Box-Only, Unpaired and Paired) at 30 min post training, g) Immunohistochemistry data for SOM tdTomato sections showing p-S6 (S235/6) in SOM neurons in CeL across groups. Scale bar, 50 μm.

**Extended Data Figure 3.**
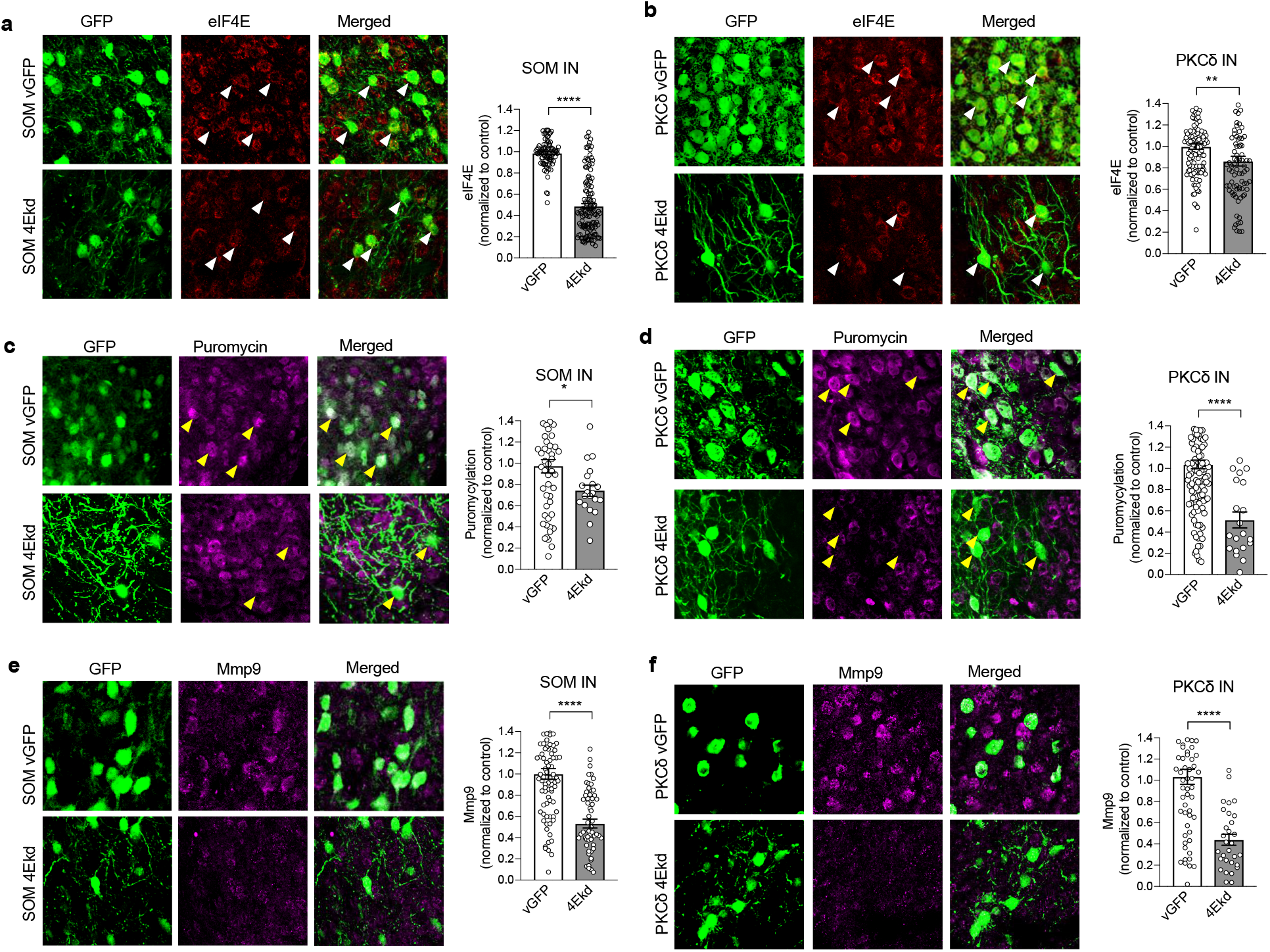
Cell type specific knockdown of cap dependent translation in CeL interneurons. a) elF4E level was significantly reduced in SOM INs in SOM 4Ekd group compared to SOM vGFP control. Unpaired t-test, p<0.0001. n=87-132/group. b) elF4E level was significantly knocked down in PKCδ INs in PKCδ 4Ekd group compared to PKCδ vGFP control. Unpaired t-test, p=0.0056, n=87-121/group. c) Global *de novo* translation, as measured with puromycin assay, was significantly reduced in SOM 4Ekd group compared to control. Unpaired t-test, p=0.0363. n=20-53/group. d) Similarly, global *de novo* protein synthesis was significantly diminished in PKCδ 4Ekd group compared to control. Unpaired t-test, p<0.0001. n=20-124/group. e) MMP9 levels was significantly reduced in SOM 4Ekd mice compared to control. Unpaired t-test, p<0.0001. n=60-87/group. f) Similarly, MMP9 level was significantly reduced in PKCδ 4Ekd group compared to control.). Unpaired t-test, p<0.0001. n=30-60/group. Data are presented as mean +SEM. *p<0.05, **p<0.01, ***p<0.001, ****p<0.0001. n.s. non significant.

**Extended Data Figure 4.**
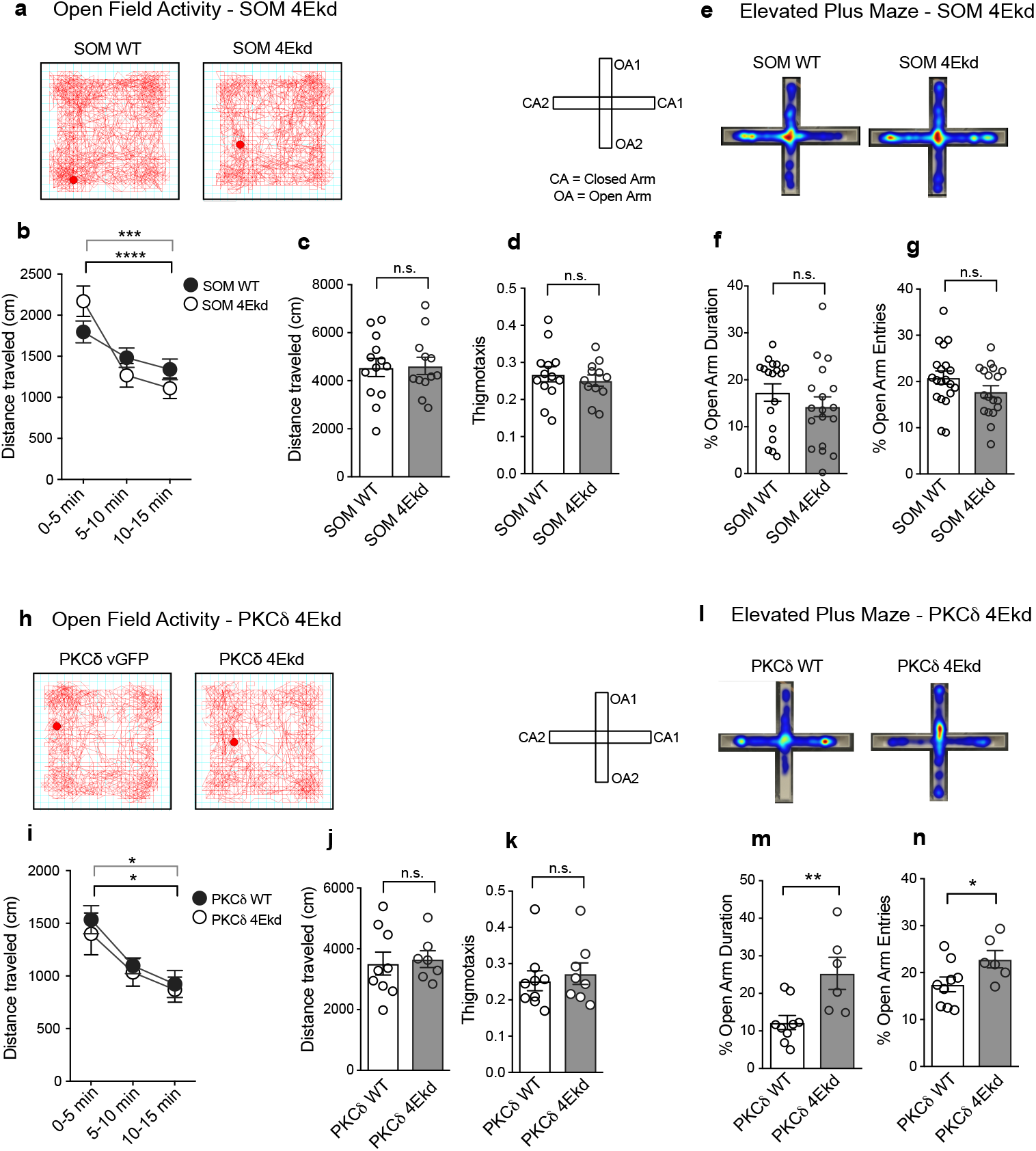
Inhibition of cap-dependent translation and anxiety related behaviors. a) Representative open field activity traces for SOM WT and SOM 4Ekd animals, b) XY plot showing normal acclimation of SOM WT and SOM 4Ekd animals to the open field arena. Interaction (Genotype X Time): F(2,46)=8.37, p=0.0008; Time: F(2,46)=45.50, p<0.0001. n=11-13/group. c) SOM WT and SOM 4Ekd animals display equivalent spontaneous locomotion in the open field arena, d) SOM 4Ekd mice display normal thigmotaxis behavior compared to control, e) Representative activity heat map in elevated plus maze for SOM WT and SOM 4Ekd animals. f) SOM WT and SOM 4Ekd animals spend similar duration in the open arm, as a percent of total duration, g) SOM WT and SOM 4Ekd mice make equivalent entries into the open arm. n=12-13/group. h) Representative open field activity traces for PKCδ WT and PKCδ 4Ekd animals, i) XY plot showing normal acclimation of PKCδ WT and PKCδ 4Ekd animals to the open field arena. Time: F(2,32)=19.12, p<0.0001. n=7-9/group. j) Bar plot showing total distance traveled by PKCδ WT and PKCδ 4Ekd mice in the open field arena, k) PKCδ 4Ekd mice show normal thigmotaxis in the open field arena compared to PKCδ WT control, n=7-9/group. I) Representative activity heat maps in elevated plus maze for PKCδ WT and PKCδ 4Ekd animals, m) Bar plot showing significantly increased %time spent in the open arm for PKCδ 4Ekd animals compared to PKCδ WT controls, p=0.0074. n=6-9/group. n) Bar plot showing % entries into the open arm, p=0.0476. n=6-9/group. Statistical tests: RM Two-way ANOVA with Bonferroni’s post-hoc test (b, i), Unpaired t-test (c, d, f, g, j, k, m, n). Data are presented as mean +SEM. **p<0.01, ****p<0.0001, n.s. non significant.

**Extended Data Figure 5.**
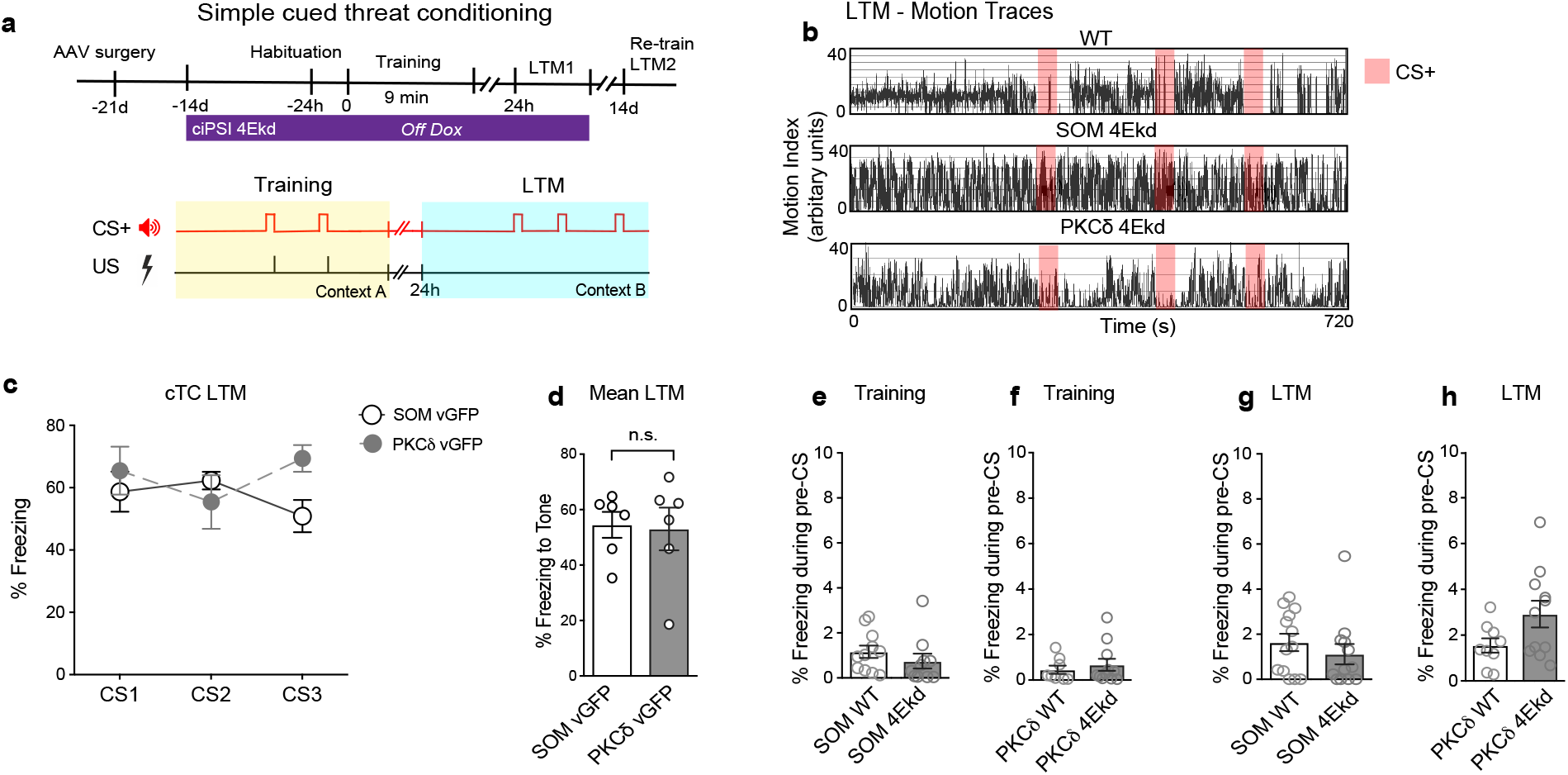
Inhibition of cap-dependent translation in CeL interneurons and simple threat conditioning. a) Schematic for simple threat conditioning paradigm in SOM and PKCδ 4Ekd mice, b) Representative motion traces for WT, SOM 4Ekd and PKCδ 4Ekd groups during LTM test, c) XY plot for SOM vGFP and PKCδ vGFP mice in simple threat conditioning showing %freezing response to successive CS presentations during LTM test (left), n=6/group. RM One-way ANOVA. Genotype: F(1,9)=0.786. d) Bar plot showing mean LTM for SOM vGFP and PKCδ vGFP mice (right). Unpaired t-test, p=0.8732. n=6/group. d) SOM 4Ekd mice and e) PKCδ 4Ekd mice have negligible freezing response during pre-CS in Training phase compared to controls, g) SOM 4Ekd mice and e) PKCδ 4Ekd mice have comparable low freezing response during pre-CS in LTM test compared to controls. RM Two-way ANOVA (c), Unpaired t-test (d-h). Data are presented as mean +SEM. **p<0.01, ****p<0.0001, n.s. non significant.

**Extended Data Figure 6.**
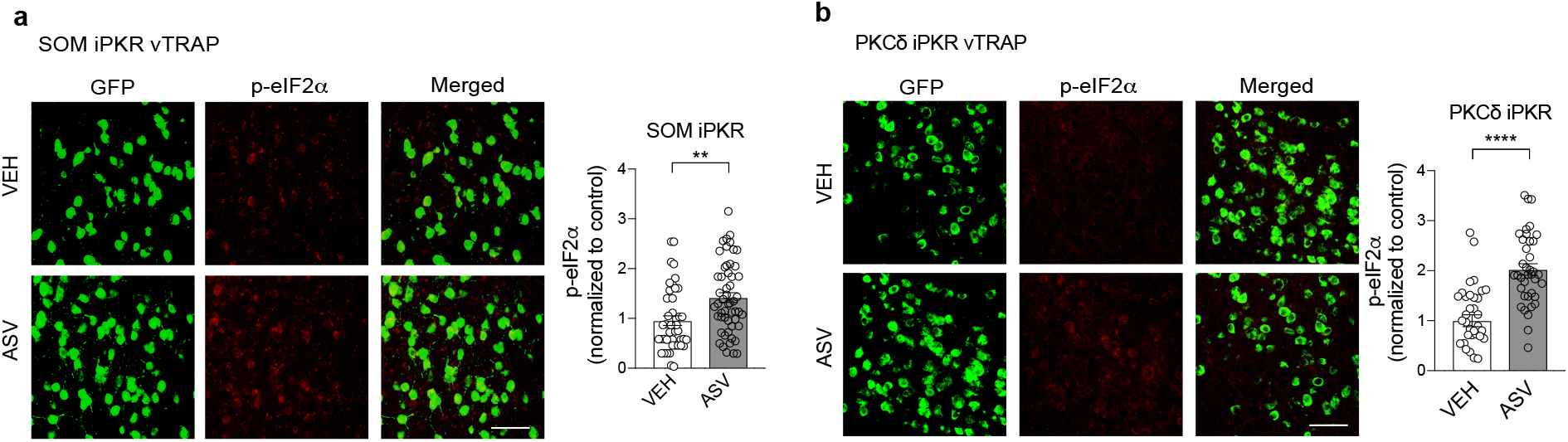
Drug-inducible cell type-specific eIF2α phosphorylation, a) Compared to vehicle controls, ASV infusion in central amygdala of SOM iPKR vTRAP animals significantly increased phosphorylation of eIF2α in SOM neurons. Unpaired t-test, p=0.0013. n=43-53/group. b) ASV infusion in CeA of PKCδ iPKR vTRAP mice also significantly elevated p-eIF2α in PKCδ neurons compared to vehicle control. Unpaired t-test, p<0.0001. n=36-38/group. c) Representative motion traces for infused WT, SOM iPKR and PKCδ iPKR mice during LTM test. All animals were infused with ASV immediately after training. CS+ in red block, CS- in green block. Data are presented as mean +SEM. **p<0.01, ****p<0.0001. Scale bar, 50 μm.

**Extended Data Figure 7.**
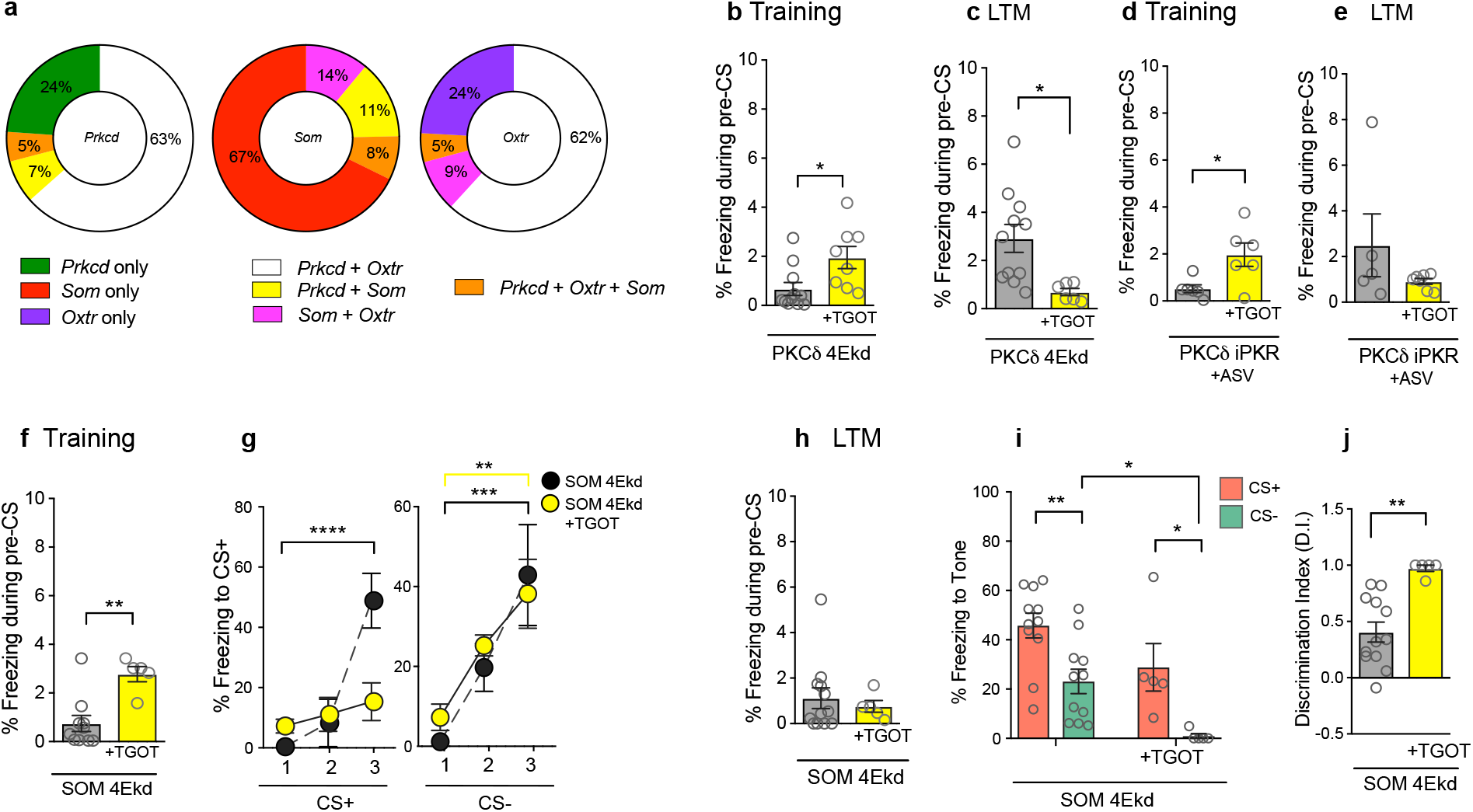
Behavioral effects of TGOT during differential threat learning. a) Co-expression analysis of *Prkcd, Som* and *Oxtr* mRNA in CeL. 63% of *Prkcd+* cells are *Oxtr+*, 62% of *Oxtr+* cells are *Prkcd+* whereas a small minority of *Som+* cells, 14%, are *Oxtr+*. b) Intra-CeL infusion of TGOT increases pre-CS freezing response in PKCδ 4Ekd animals during training. Unpaired t-test, p=0.019, n=8-11/group. c) Pre-training TGOT infusion leads to a significant decrease in freezing response during pre-CS of LTM test. Unpaired t-test, p=0.0144, n=6-11/group. d) Intra-CeL infusion of TGOT also increases pre-CS freezing response in PKCδ iPKR +ASV mice. n=6/group e) Pre-training TGOT infusion does not alter freezing response during pre-CS of LTM test. g) Pre-training infusion of TGOT in CeA of SOM 4Ekd mice impaired the learning curve for CS+ during training (left). RM Two-way ANOVA with Bonferroni’s post-hoc test. Interaction (Drug X CS+): F(2,16)=12.29, p=0.0006. n=5/group. However, TGOT did not affect successive freezing response to CS- in SOM 4Ekd mice. RM Two-way ANOVA with Bonferroni’s post-hoc test. CS-: F(2,16)=19.23, p<0.0001. n=5/group. h) TGOT has no effect on freezing response in SOM 4Ekd mice during pre-CS of LTM test. i) TGOT significantly decreases conditioned safety response to CS- in SOM 4Ekd mice. Two-way ANOVA with Bonferroni’s post-hoc test. Drug: F(1,28)=9.935, p=0.0038, CS: F(1,28)=16.61, p=0.0003. n=5-12/group. j) Discrimination index is highly enhanced for SOM 4Ekd mice following TGOT infusion. Unpaired t-test, p=0.0012. n=5-12/group. Data are presented as mean +SEM. *p<0.05, **p<0.01, ***p<0.001, ****p<0.0001. n.s. non significant.

**Extended Data Figure 8.**
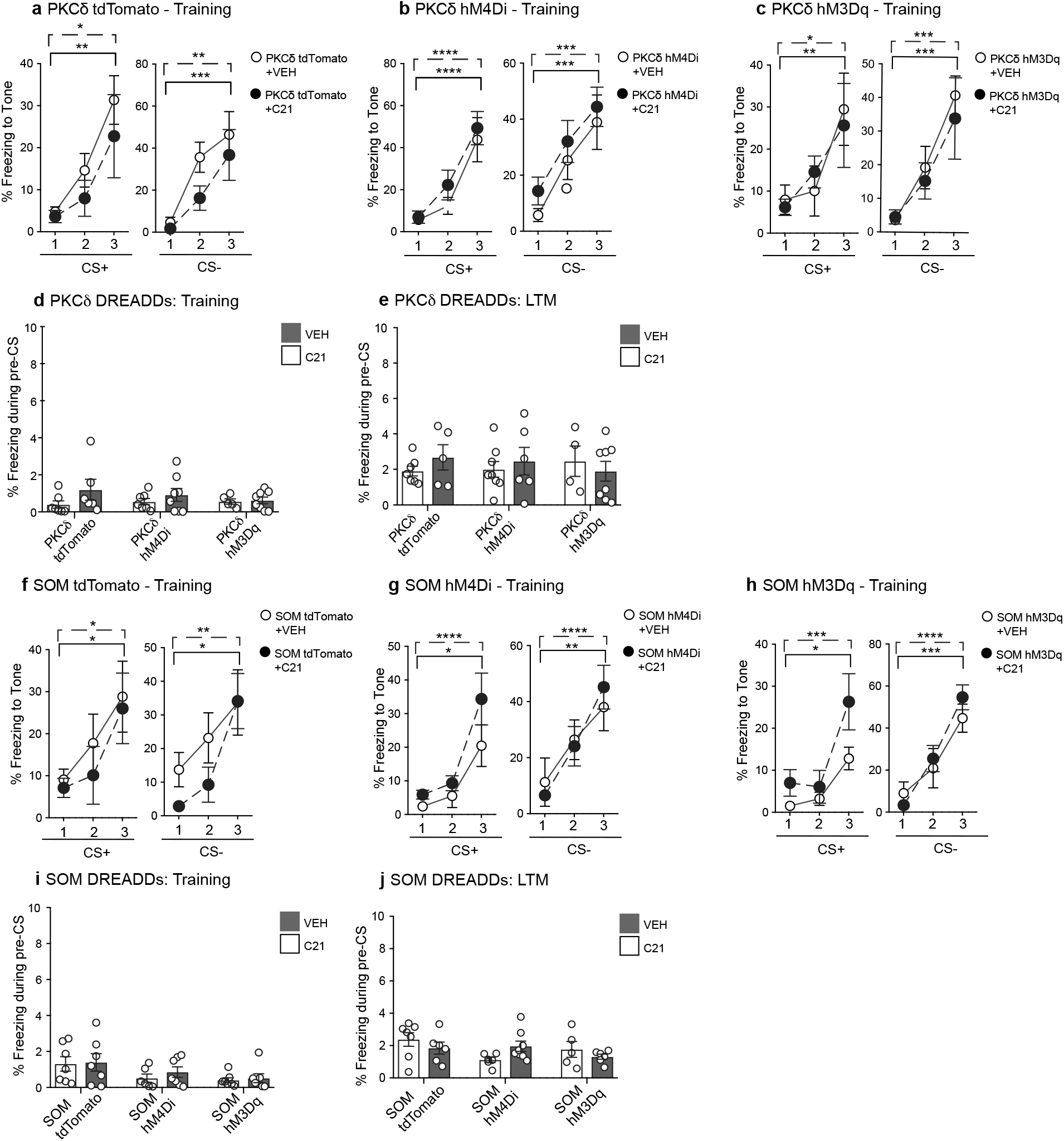
Chemogenetic modulation of G-protein signaling in CeL interneurons affects associative learning. a) C21 treated PKCδ tdTomato animals have normal memory acquisition relative to VEH controls, with progressive increase in freezing response to successive presentation of CS’s. CS+: F(2,22)=11.96, p=0.0003; CS-: F(2,22)=20.01, p<0.0001. n=7/group. b) C21 treated PKCδ hM4Di animals learn normally compared to VEH controls. F(2,26)=32.94, p<0.0001; CS-: F(2,26)=32.94, p<0.0001. n=7-9/group. c) C21 treated PKCδ hM3Dq animals acquire differential threat memory normally compared to VEH controls. CS+: F(2,24)=12.20, p=0.0002; CS-: F(2,24)=24.62, p<0.0001. n=7/group. d) Freezing response during pre-CS of Training phase is negligible across all C21 and VEH treated PKCδ groups. e) C21 treatment in PKCδ tdTomato, PKCδ hM4Di and PKCδ hM3Dq animals does not alter baseline freezing response during pre-CS of LTM test. f) C21 treated SOM tdTomato mice learn normally compared to VEH treated controls. CS+: F(2,22)=8.02, p=0.0024; CS-: F(2,22)=17.00, p<0.0001. n=6-7/group. g) C21 treated SOM hM4Di mice have normal memory acquisition relative to VEH controls. CS+: F(2,22)=20.62, p<0.0001; CS-: F(2,22)=19.62, p<0.0001. n=6-7/group. h) C21 treated SOM hM3Dq animals acquire differential threat memory normally relative to VEH controls. CS+: F(2,20)=17.09, p<0.0001; CS-: F(2,20)=38.94, p<0.0001. n=5-7/group. i) Freezing response during pre-CS of Training phase is negligible across all C21 and VEH treated SOM groups. j) C21 treated SOM tdTomato, SOM hM4Di and SOM hM3Dq mice have equivalent freezing response during pre-CS compared to VEH controls. Statistics: RM Two-way ANOVA with Bonferroni’s post-hoc test (a-f). Data are presented as mean +SEM. *p<0.05, **p<0.01, ***p<0.001, ****p<0.0001. n.s. non significant.

**Extended Data Figure 9.**
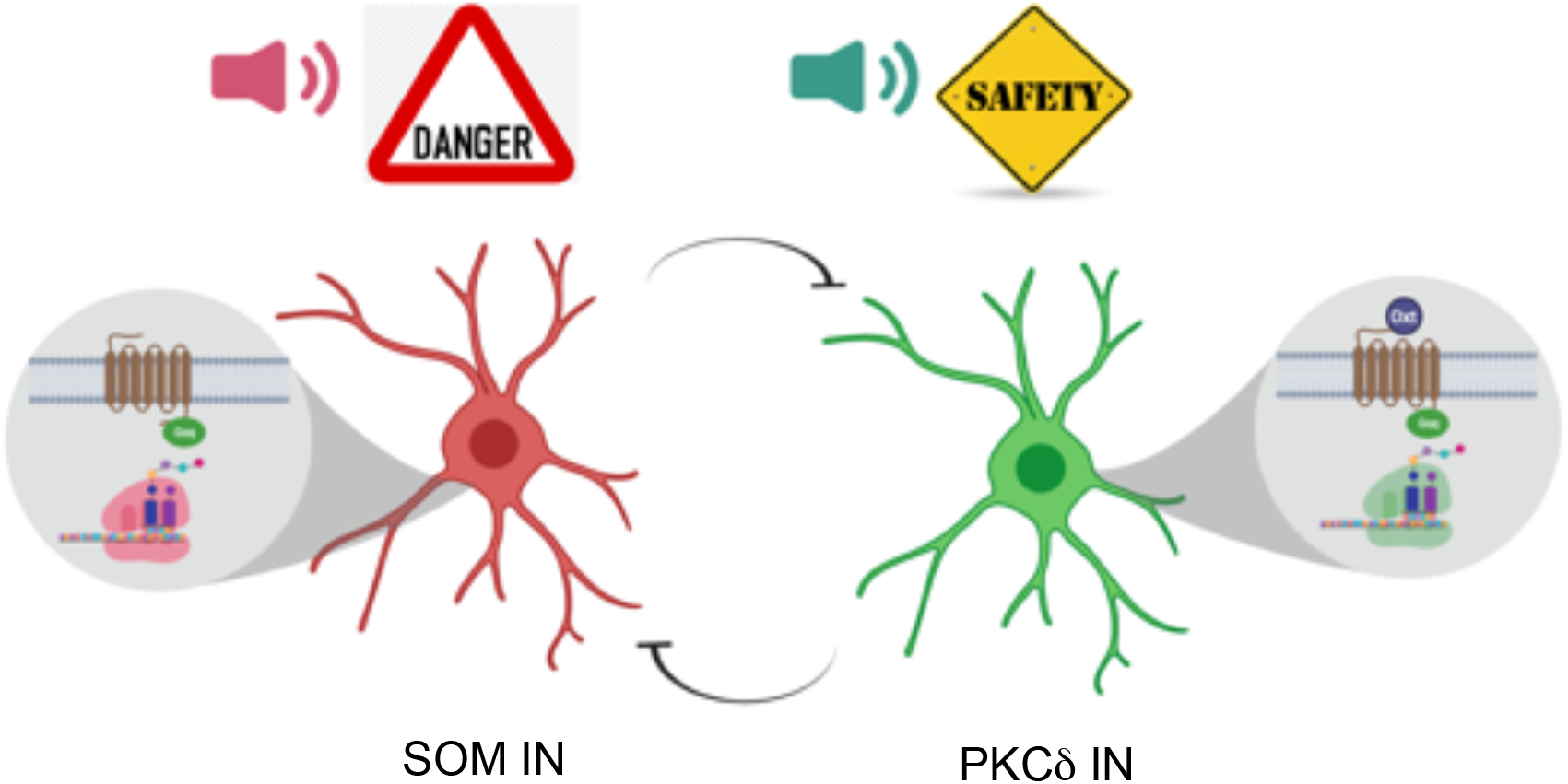
Working model of simultaneous storage of threat and safety cue associated memories in CeL SOM and PKCδ INs respectively.

